# Molecular Basis of Histone H3 Reading and Writing by *Legionella pneumophila* SET Domain Lysine Methyltransferases

**DOI:** 10.64898/2026.05.29.728679

**Authors:** Stephen Gonzalez, Thomas Knight, Juan M. Nevarez, Alexandra H. Evans, Kayla M. Daniels, Emily Xu, Anup Vaidya, Jonathan M. Burg, Oluwatobi A. Adeleke, Shahrad Daraeikia, Saarang Gopinath, Michaela Yamine, Matthew Yacoub, Sojin An, Ethan Madison, Jean-Marc Fontaine, Michele Swanson, Michael-Christopher Keogh, Raymond C. Trievel

## Abstract

RomA and its highly conserved strain ortholog LegAS4 are SET and ankyrin domain-containing effector proteins of the intracellular bacterial pathogen *Legionella pneumophila*. These enzymes are secreted into host cells, where they translocate to the nucleus and methylate Lys14 in histone H3 (H3K14), a novel post-translation modification (PTM) that reprograms gene expression and promotes bacterial replication. To elucidate their H3K14 substrate specificity, we determined the crystal structures of LegAS4 and RomA bound to histone H3 peptides and characterized nucleosome binding and methylation by the enzymes. The results reveal a distinctive bipartite engagement of the histone H3 N-terminal tail by the enzymes’ SET and ankyrin domains. Further, the ankyrin domain distinguishes different PTMs at H3R2 and H3K4, which is crucial for nucleosome engagement and H3K14 methylation. Together, these studies yield new insights into histone H3 reading and writing by the *L. pneumophila* histone lysine methyltransferases during host cell infection.

## Introduction

*Legionella pneumophila* is the primary causative agent of Legionnaires’ Disease, a bacterial pneumonia that primarily afflicts the elderly or individuals who are immunocompromised or have chronic lung disease^1, 2, 3^. This environmental gram-negative bacterium is a facultative intracellular pathogen whose primary hosts include various species of amoeba and other phagocytic protozoa inhabiting natural and engineered freshwater systems and soils. Humans contract Legionnaires’ Disease after inhaling aerosolized water droplets containing *L. pneumophila* into the lungs, where the pathogen replicates within alveolar lung macrophages.

*L. pneumophila* initiates infection by secreting >300 different proteins into host phagocytes^4, 5^. Particular effectors promote pathogen survival and replication by subverting a diverse array of host cellular pathways, including metabolism, intracellular trafficking, signal transduction, autophagy, apoptosis, and gene expression. Among the list of characterized *L. pneumophila* nuclear effectors are histone lysine methyltransferases (LpHKMTs) and histone deacetylases^6, 7, 8, 9^. After secretion into the host cell, these enzymes translocate to the nucleus where they function to alter chromatin modifications. These changes reprogram host gene transcription and promote bacterial replication in part by dampening the expression of innate immune response genes^6, 9^.

The LpHKMTs currently comprise two enzymes: Regulation of Methylation A (RomA) and its highly conserved ortholog LegAS4 from the *L. pneumophila* Paris and Philadelphia strains, respectively^6, 7^. The protein architecture of the LpHKMTs comprises an N-terminal lysine-rich nuclear localization signal (NLS); a Su(var)3-9, Enhancer-of-zeste, and Trithorax (SET) domain that catalyzes S-adenosylmethionine (AdoMet)-dependent lysine methylation; and a C-terminal ankyrin repeat-containing domain^6, 7^. RomA specifically methylates Lys14 in histone H3 (H3K14), a novel post-translational modification (PTM) not observed in mammalian chromatin^6^. During host cell infection, RomA localizes to the nucleus and catalyzes global H3K14 methylation, thereby repressing innate immunity genes, including the pro-inflammatory cytokine interleukin-6 and Toll-like receptor 5 (TLR5) that initiates an immune signaling cascade upon recognition of bacterial flagellin^10^. Deletion of the *romA* gene in *L. pneumophila* strain Paris impairs intracellular replication in infected host cells compared to the wild-type (WT) strain, illustrating its importance in pathogenesis^6^.

LegAS4 remains less characterized than RomA. Initial studies indicated that LegAS4 selectively methylates H3K4 and to a lesser degree H3K9 *in vitro* and during host cell infection localizes to the nucleoli, where it stimulates rDNA transcription^7, 11^. Such findings starkly contrast with the H3K14 specificity and transcriptional repression mediated by RomA, leading to speculation that the two LpHKMTs possess different substrates and pathogenic functions due to sequence variations in their N-termini^12^. However, more recent structural and functional studies have shown that LegAS4 recognizes and methylates H3K14 *in vitro* and mediates global H3K14 methylation in the nucleus of host cells infected by *L. pneumophila* strain Philadelphia-1^13, 14^. Thus, RomA and LegAS4 share a conserved histone lysine specificity and role in *Legionella* pathogenesis.

To elucidate the substrate specificity and molecular functions of LegAS4 and RomA, we characterized their nucleosome engagement, methyltransferase activity, and crystal structures in complex with histone H3 N-terminal tail peptides. Our biochemical studies reveal that LegAS4 and RomA bind nucleosomes with apparent nanomolar affinity and specifically methylate H3K14. In agreement with these observations, the structures of LegAS4 and RomA bound to H3 peptides illustrate that the histone residues flanking H3K14 mediate a conserved network of interactions with invariant residues in the enzymes’ SET domains. Further, the ankyrin repeat domain of the LpHKMTs binds to the N-terminal residues in histone H3, recognizing the PTM status of H3R2 and H3K4 to regulate nucleosome association and H3K14 methylation. These results furnish novel insights into the molecular mechanisms of nucleosome engagement and histone H3 reading and writing by the LpHKMTs.

## Results

### LegAS4 methylates H3K14 in host cells and *in vitro*

To resolve the histone lysine specificity of LegAS4, we examined its activity in infected host cells. To do so, we generated a deletion of the *legAS4* gene (*ΔlegAS4)* in the Lp02 strain, a thymine auxotroph derived from *L. pneumophila* Philadelphia-1^15^. Mouse bone marrow-derived macrophages (BMDMs) were infected with either WT Lp02, the Lp02 *ΔlegAS4* mutant, or the *ΔlegAS4* mutant transformed with an empty vector control. Immunoblot analysis from 1 – 6 hours post-infection revealed increasing H3K14 methylation in the BMDMs infected by WT Lp02 but not those infected by the Lp02 *ΔlegAS4* mutant or its empty vector transformant (Figure 1A, S1A, S2A). A similar time course of H3K14 methylation was observed in host cells infected by the *L. pneumophila* Paris strain^6^. To assess if H3K14 methylation could be restored in Lp02 *ΔlegAS4*, we performed complementation studies with either WT *legAS4*, WT *romA* from *L. pneumophila* Paris, or a catalytically defective (CD) *legAS4* allele harboring six mutations in the AdoMet binding cleft and active site (Tyr135, Ser158, Tyr159, Asn181, Tyr220, and Tyr231; LegAS4 amino acid numbering reflects a recently reported alternative translation start site^14^) (Figure S1B, S1C). Each gene was integrated in the Lp02 genome at a neutral site that does not perturb the expression of neighboring genes^16^. Immunoblot analysis of infected BMDMs illustrated that H3K14 methylation was restored in Lp02 *ΔlegAS4* encoding either *legAS4* or *romA* but not *legAS4* CD (Figure 1B, S2B). Together, these findings demonstrate that *legAS4* is responsible for H3K14 methylation in cells infected by the Lp02 strain of *L. pneumophila*.

**Figure 1:**
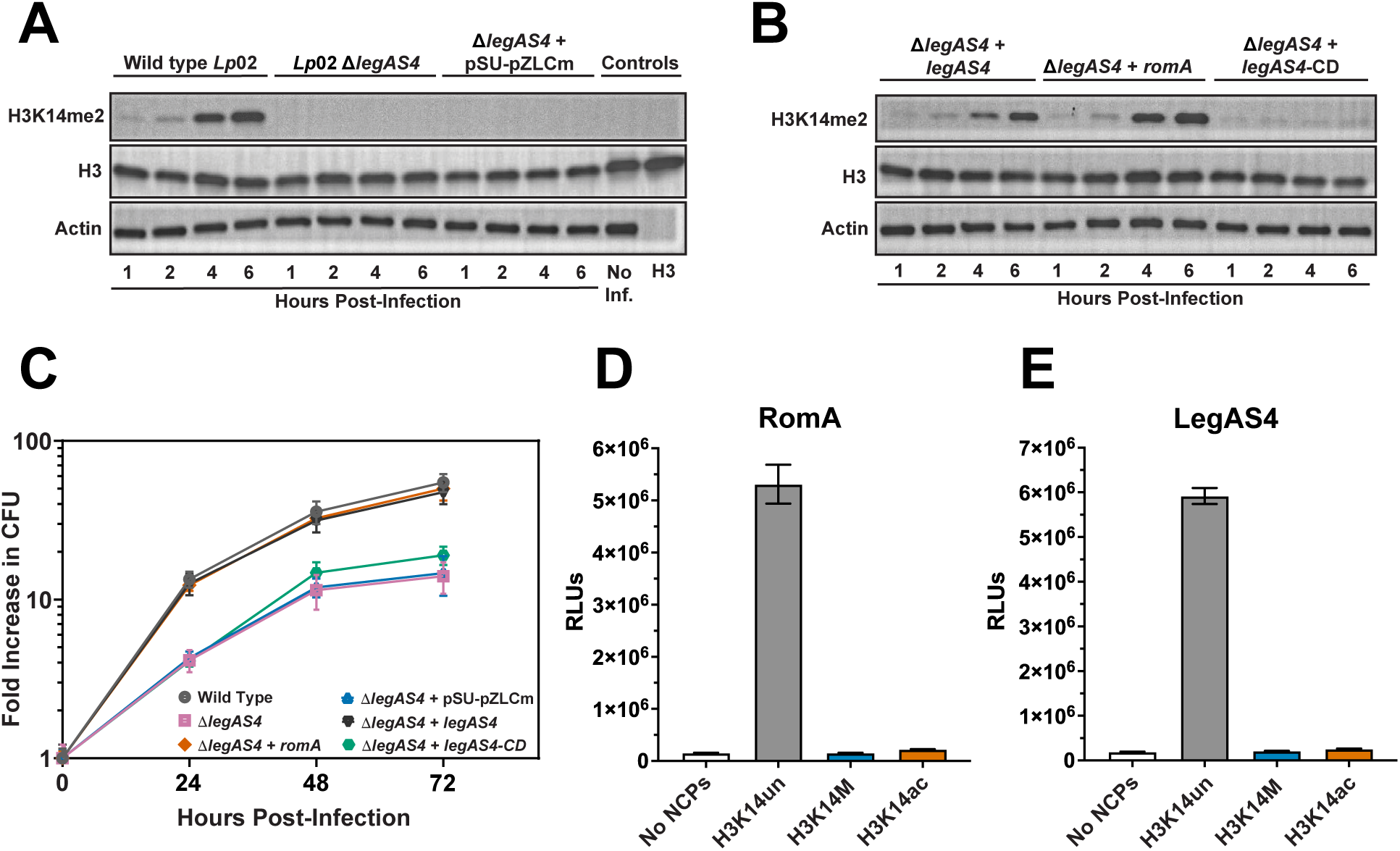
LegAS4 methylates H3K14 in cells and *in vitro*: **A.** Western blots of H3K14 methylation in mouse bone marrow-derived macrophages (BMDMs) infected with *L. pneumophila* wild-type (WT) strain Lp02, Δ*legAS4* mutant, or Δ*legAS4* mutant transformed with control plasmid pSU-pZLCm. H3K14 methylation was monitored from 1 – 6 hours post-infection and in uninfected (No Inf.) macrophages. Histone H3 and β-actin served as loading controls. Anti-H3K14me2 was validated to PTM-defined nucleosomes (Figure S1A). **B.** Complementation assay western blots in which a *legAS4*, *romA*, or *legAS4*-catalytically defective (CD) mutant gene was integrated to a neutral genomic site of the Lp02 Δ*legAS4* mutant genome. Histone H3 and β-actin were loading controls as in (**A**). **C.** Replication in BMDMs of WT Lp02, the Lp02 Δ*legAS4* mutant, Δ*legAS4+*pSU-pZLCm control plasmid, Δ*legAS4+legAS4,* Δ*legAS4+romA*, and Δ*legAS4*+*legAS4*-catalytically defective (CD) mutant. Colony forming units (CFUs) were quantified over a 72-hour period. **D,E.** *In vitro* luminescent methyltransferase assays of RomA (**D**) and LegAS4 (**E**) using recombinant nucleosome core particles (NCPs), including *Xenopus laevis* ([H3.1]_2_), a H3K14 to methionine substitution ([H3.1K14M]_2_), and human ([H3K14ac]_2_) NCPs (Table S1). As negative controls, assays were also performed with no NCPs. Activity was measured in relative luminescent units (RLUs). Assays were performed in triplicate, and error bars represent standard deviation of the mean.

To evaluate the impact of methylation by LegAS4, we quantified intracellular replication by the WT Lp02 strain, the Lp02 *ΔlegAS4* mutant, and its *ΔlegAS4* genetic complements. Lp02 *ΔlegAS4* exhibited impaired replication in BMDMs compared to WT Lp02 (Figure 1C, S2C), consistent with the diminished replication of the *ΔromA* mutant of the *L. pneumophila* Paris strain in either protozoan *Acanthamoeba castellanii* or monocytic THP-1 cells^6^. Complementation of *ΔlegAS4* with *legAS4* or *romA* restored intracellular replication comparable to WT Lp02, whereas *ΔlegAS4* transformed with the empty vector control or *legAS4* CD did not (Figure 1C). Thus, LegAS4 promotes efficient replication of WT Lp02 in BMDMs, and *romA* can genetically complement the *ΔlegAS4* mutant.

Our observation that LegAS4 catalyzed H3K14 methylation during macrophage infection prompted us to examine the substate specificity of purified recombinant LpHKMTs towards nucleosomes *in vitro*. In luminescent methyltransferase assays, RomA and LegAS4 each displayed robust activity toward recombinant unmodified nucleosomes (Figure 1D, 1E, S2D, S2E and Table S1). In contrast, methylation by the enzymes was abolished in homotypic nucleosomes bearing either an H3K14 to methionine mutation ([H3.1K14M]_2_) or H3K14 acetylation ([H3K14ac]_2_) (here we use a recently proposed nomenclature for improved communication in chromatin studies^17^: see Methods). These *in vitro* activities are consistent with the *legAS4*-dependent methylation of H3K14 observed in Lp02-infected BMDMs (Figure 1A, 1B) and studies illustrating H3K14 methylation by RomA and LegAS4 in host cells respectively infected by the *L. pneumophila* Paris and Philadelphia-1 strains^6, 13, 14^. In summary, our *in vitro* and cell-based assays demonstrate that the LpHKMTs are H3K14-specific methyltransferases, and abrogating LegAS4 methylation of H3K14 impairs the intracellular replication of Lp02 in macrophages.

### Nucleosome engagement by the LpHKMTs

Our findings that RomA and LegAS4 robustly and site specifically methylate nucleosomes spurred us to investigate the mechanism of enzyme–substrate recognition. To accomplish this aim, we utilized Captify™ amplified luminescent proximity homogeneous assays (Captify-Alpha) to analyze the binding of WT RomA or a truncated form lacking its N-terminal NLS (RomA βNLS) to unmodified nucleosomes or a 147 base-pair 601 DNA (Figure 2A, S1C, and Table S1, S2)^18, 19^. WT RomA bound to nucleosomes at nanomolar concentrations but displayed weaker binding to the free 601 DNA, indicating that both histone and DNA interactions mediate enzyme–nucleosome engagement. Notably, RomA βNLS exhibited dramatically reduced binding to nucleosomes and free 601 DNA, presumably due to the loss of electrostatic interactions between the basic NLS motif and the DNA phosphate backbone (Figure 2A, S1C). Corroborating these observations, RomA and LegAS4 bound nucleosomes with comparable affinity in electrophoretic mobility shift assays (EMSAs), whereas nucleosome recognition by RomA βNLS was undetectable (Figure 2B, 2C, S3A, S3B). Together, these findings reveal that the NLS motif of the LpHKMTs exhibits dual functions in facilitating not only nuclear localization in the host cell^6, 7^ but also nucleosome engagement.

**Figure 2:**
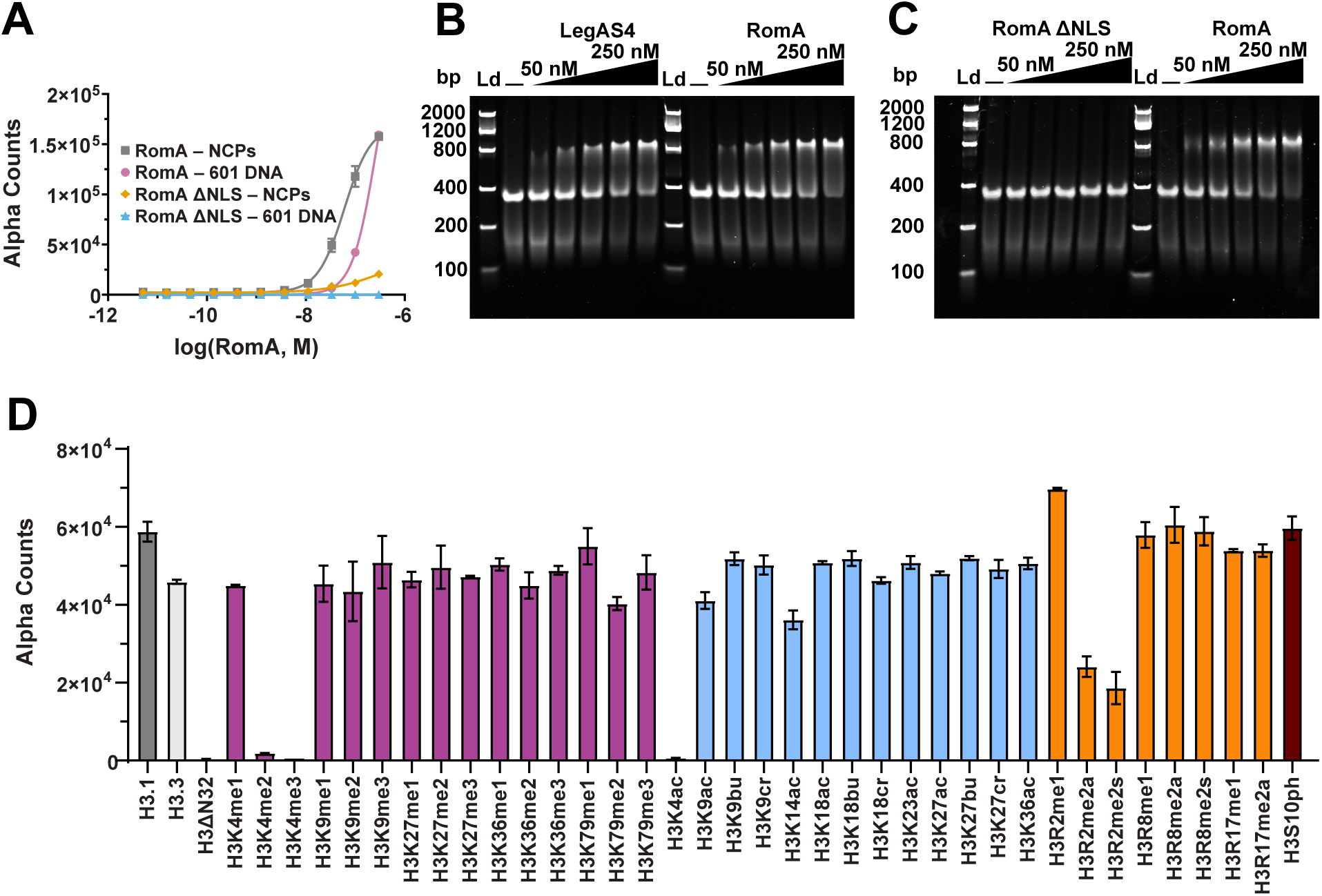
Nucleosome engagement by RomA. **A.** Captify™-Alpha was used to measure the binding of WT RomA and RomA ΔNLS (titrated) to human ([H3.1]_2_) NCPs or 147 bp 601 DNA oligonucleotide (constant). Relative binding units were expressed in amplified luminescence proximity homogeneous assay (Alpha) counts. **B.** EMSAs using increasing concentrations of LegAS4 or RomA with a fixed concentration of recombinant *X. laevis* ([H3.1]_2_) NCPs (25 nM). **C.** EMSAs of WT RomA and RomA ΔNLS with ([H3.1]_2_) performed as in (**B**). **D.** Captify-Alpha was used to profile the association of WT RomA with a panel of fully-defined human NCPs bearing histone H3 variants (([H3.1]_2_), ([H3.3]_2_)), a tail-deletion ([H3ΔN32]_2_), and H3 bearing distinct PTMs, including lysine methylation (violet), acylation (light blue), arginine methylation (orange), and phosphorylation (dark red) (Table S1).

Having established that the LpHKMTs directly bind nucleosomes, we next investigated the impact of histone context. Utilizing Captify-Alpha, we profiled RomA binding to a panel of defined nucleosomes bearing histone variants, N-terminal tail truncations, or PTMs (Table S1). The profiling illustrated that none of the tested histone variants (e.g., histone H3.1, H3.3, H2AX, H2AZ.1, and H2AZ.2) had an appreciable impact on nucleosome binding by RomA (Figure 2D, S3C). In contrast, RomA engagement was abolished by a nucleosome bearing a truncation of histone H3 residues 1 – 32 (termed ([H3βN32]_2_)) that harbors the H3K14 methylation site, identifying this region as essential for enzyme binding (Figure 2D). On the other hand, deletion of residues 1 – 15 in the histone H4 N-terminal tail, as in ([H4ΔN15]_2_), did not perturb nucleosome association (Figure S3D).

Specific PTMs in the core histones differentially impacted nucleosome binding by RomA. Modifications in histones H2A, H2B, H4, and most of H3 had no appreciable effects on nucleosome recognition, in stark contrast to specific PTMs at H3R2 and H3K4 (Figure 2D, S3C, S3D). Notably, ([H3R2me1]_2_) slightly increased RomA binding, whereas ([H3R2me2a]_2_) and ([H3R2me2s]_2_) modestly diminished enzyme association. In contrast, H3K4 PTMs had a more pronounced impact on nucleosome engagement. Nucleosomes with unmodified H3K4 (termed ([H3K4un]_2_)) and ([H3K4me1]_2_) were similarly engaged by RomA, whereas binding was abolished to ([H3K4me2]_2_), ([H3K4me3]_2_), and ([H3K4ac]_2_). Together, these findings suggest the LpHKMTs can distinguish the PTM landscape at H3K4 and, to a lesser extent, at H3R2.

### Structural basis of histone H3 recognition by LegAS4 and RomA

To elucidate the histone substrate specificity of the LpHKMTs, we determined a 1.6 Å resolution crystal structure of a LegAS4 lacking its NLS motif in complex with the methyl transfer product S-adenosylhomocysteine (AdoHcy) and a 21-mer histone H3 peptide (H3_[1-21]_) (Figure 3A and Table S3, S4). The LegAS4 structure adopts a “V” shape with the SET and ankyrin domains each forming one arm. The refined structure reveals electron density for histone H3 residues 1 – 18 that nestle in the nook of the V and span the two domains. Strong electron density is observed across the histone H3 main and side chains, with two exceptions: 1) the side chain of H3R17 that adopts alternate conformations and 2) the side chain of H3K18 that is disordered and thus was not modeled (Figure S4A – S4C). Notably, the binding mode observed for our structure bound to the H3_[1-21]_ peptide differs from a previously determined structure of LegAS4 bound to a histone H3 peptide comprising residues 3 – 17 (H3_[3-17]_) ((Figure S4D)^14^. Truncation of H3A1 and H3R2 in the H3_[3-17]_ peptide induces residues H3T3 – H3T11 to adopt out of register conformations compared to the structure of LegAS4 bound to the H3_1-21_ peptide. We propose that the unambiguous electron density observed for H3A1 and H3R2 coupled with their direct interactions with the enzyme confirm that the LegAS4/AdoHcy/H3_[1-21]_ peptide complex represents the physiologically relevant binding mode for histone H3.

**Figure 3:**
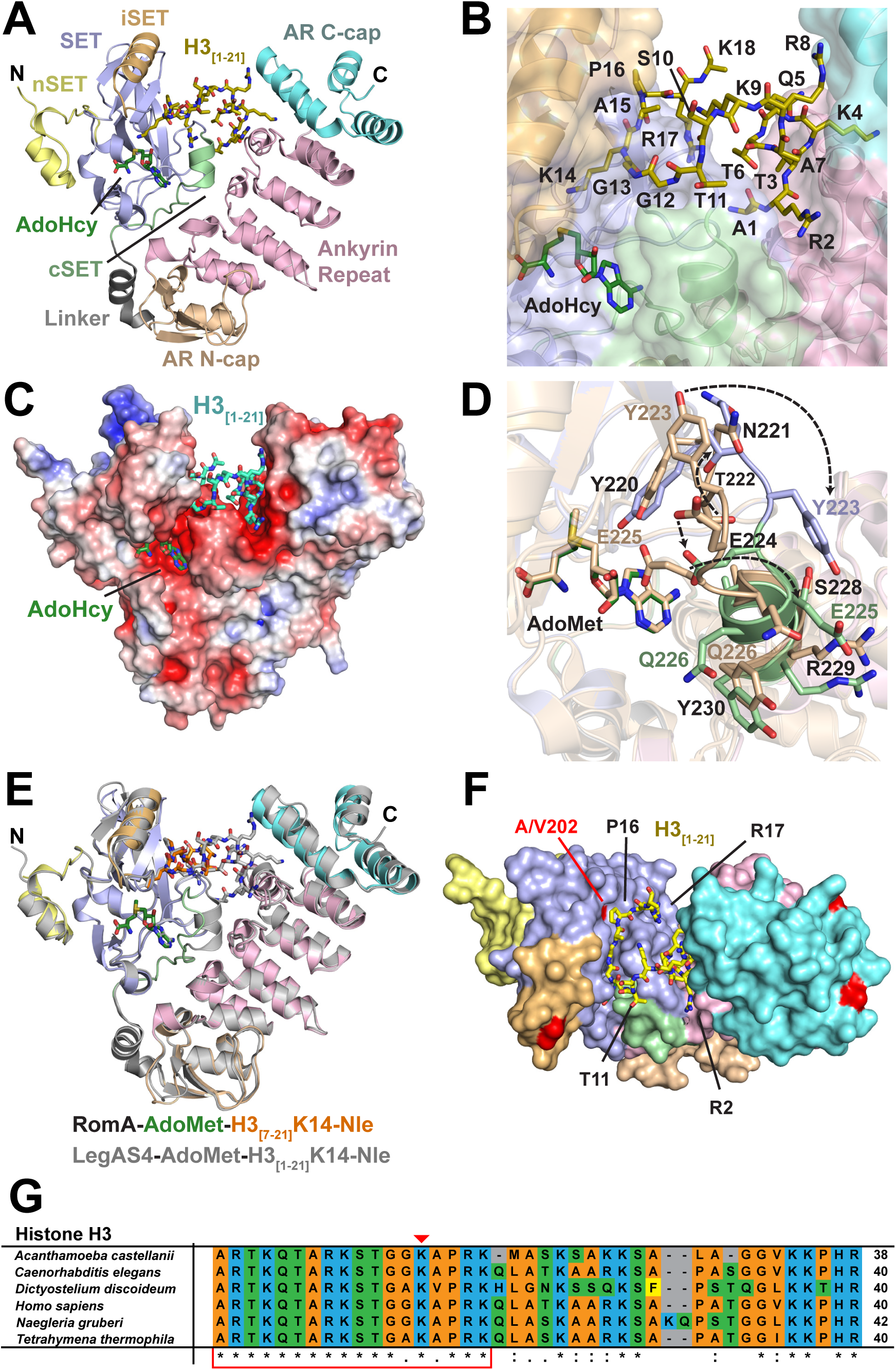
Recognition of histone H3 N-terminal tail by LegAS4 and RomA. **A.** Cartoon renderings of the 1.6 Å resolution crystal structure of LegAS4 bound to product S-adenosylhomocysteine (AdoHcy) and a histone H3_[1-21]_ peptide. The various domains and subdomains in LegAS4 are denoted by the corresponding colors in the LpHKMT sequence alignment (Figure S1C). The H3_[1-21]_ peptide and AdoHcy are depicted with gold and green carbon atoms, respectively. **B.** Semi-transparent surface of LegAS4 illustrates the bipartite engagement of the H3_[1-21]_ peptide residues by the SET and Ankyrin repeat domains. **C.** Electrostatic surface of the LegAS4/AdoHcy/H3_[1-21]_ peptide ternary complex. Surface potential is colored from red (−5 k_B_T/e) to blue (+5 k_B_T/e); histone H3_[1-21]_ peptide, cyan carbon atoms; AdoHcy, green carbon atoms. **D.** Superimposition of the switch region of the histone H3 binding cleft in the SET domain and cSET motif from the LegAS4/AdoMet binary complex (colored in wheat, PDB 5CZY) and LegAS4/AdoMet/H3_[1-21]_K14-Nle peptide ternary complex (colored as in **A**). Dashed arrows denote residues in the binary and ternary states that undergo large conformational changes on association with histone H3. Supplemental Movie S1 illustrates the conformational change in the switch region during histone H3 binding. **E.** Structural alignment of the RomA/AdoMet/H3_[7-21]_K14-Nle peptide (enzyme colored as in **A** with the histone H3 peptide depicted with orange carbon atoms) and LegAS4/AdoMet/H3_[1-21]_K14-Nle peptide (gray) complexes. **F.** Molecular surface representation of the LegAS4/AdoMet/H3_[1-21]_K14-Nle peptide complex, colored as in (**A**), with sequence variations between LegAS4 and RomA highlighted in red on the surface. Select histone H3 residues are denoted. **G.** Sequence alignment of the N-terminal tails of histone H3 homologs from representative protozoan and metazoan hosts of *L. pneumophila*. The text background colors signify side chain properties: aromatic (yellow), basic (blue), aliphatic (orange), and polar (green). Sequence identity (*), strong sequence conservation (:), and weak conservation (.) are indicated under the sequence alignment. Red triangle 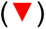 denotes the K14 methylation site, and the red box demarcates the histone H3 sequence motif recognized by the LpHKMTs.

The domain architecture of the LpHKMTs engenders a histone H3 recognition mode distinct from characterized eukaryotic SET domain KMTs^20^. In the LegAS4 ternary complex, the ankyrin and SET domains recognize the histone H3 N-terminal tail through a bipartite binding mechanism (Figure 3B). Residues H3A1 – H3R8 primarily interact with the ankyrin domain, with additional contributions from residues in the SET domain at the domains’ interface. In contrast, residues H3K9 – H3R17, which include the H3K14 methylation site, bind exclusively to the catalytic SET domain. The H3K14 side chain is deposited within the lysine binding channel that extends to the AdoMet binding pocket on the opposite face of the SET domain, as observed in the structures of other SET KMTs^21^. An extensive network comprising > 20 hydrogen bonds between histone H3 and both enzyme domains facilitates substrate recognition (Table S5). Further, the surfaces of the histone H3 and AdoMet binding sites in LegAS4 are highly acidic, promoting electrostatic interactions between the enzyme and its positively charged substrates (Figure 3C). Utilizing isothermal titration calorimetry, we measured equilibrium dissociation constants of 424 nM and 19.7 nM for AdoMet and the histone H3_[1-21]_ peptide, respectively, demonstrating that hydrogen bonding and electrostatic interactions mediate high affinity substrate binding to the LpHKMTs (Figure S5).

The extensive interactions between histone H3 and the SET and ankyrin domains prompted us to examine if H3 binding induces conformational changes in the enzyme. To this end, we determined a 1.8 Å resolution X-ray structure of a LegAS4 ternary complex bound to AdoMet and a histone H3 peptide with H3K14 substituted by a non-substrate norleucine (H3_[1-21]_K14-Nle) for comparison to a previously determined LegAS4/AdoMet binary complex^22^ (Table S3, S4). Superimposition of the LegAS4 binary and ternary complexes revealed high structural homology with a root mean squared deviation (RMSD) of 0.62 Å for all aligned Cα atoms. However, the loop at the SET domain C-terminus (residues 220 – 223) and the α-helix in the cSET region (residues 224 – 230), which we collectively term the switch region, undergo substantial rearrangements upon binding to the histone H3 N-terminal tail (Figure 3D and Movie S1). In the LegAS4 binary complex, the C-terminal loop in the switch region partially occupies the histone H3 binding cleft, analogous to the autoinhibitory loops described in the catalytic domains of certain SET domain KMTs^23, 24, 25, 26^. In the histone H3 peptide-bound ternary complex, the C-terminal loop of the switch region is shifted toward the enzyme’s ankyrin domain to accommodate substrate association with the SET domain. Several of the residues in this loop, including Asn221, Thr222, and Tyr223, interact with histone H3.

We next investigated whether the histone H3 binding mode observed for LegAS4 is conserved in RomA. Attempts to determine the structure of a RomA construct lacking its NLS motif in complex with the H3_[1-21]_ peptide were unsuccessful, but we were able to solve a 1.75 Å resolution X-ray structure of the enzyme bound to AdoMet and a 15-mer H3K14-Nle peptide comprising residues 7-21 (H3_[7-21]_K14-Nle) (Table S3, S4). Structural superimposition illustrates that LegAS4 and RomA are highly homologous with an RMSD of 0.48 Å (for all aligned Cα atoms), concurring with their high degree of sequence identity (Figure 3E, S1C). Mapping of the non-identical amino acids between RomA and LegAS4 onto the surface of LegAS4 reveals that the residues in the ankyrin and SET domains that interact with histone H3 are invariant, with the exception of Ala202 in the H3P16 binding cleft of LegAS4 which is substituted by Val202 in RomA (Figure 3F). Together, the RomA and LegAS4 structures reveal that the enzymes share a highly conserved histone H3 binding site.

The high degree of sequence conservation within the substrate binding clefts of LpHKMTs suggested that the enzymes may recognize a conserved target motif in the histone H3 N-terminal tail. As the peptides co-crystallized with LegAS4 and RomA were based on the sequence of human histone H3, we aligned the H3 N-terminal tail sequences from several protozoan and metazoan eukaryotic hosts of *L. pneumophila*, including *Acanthamoeba castellanii*, *Caenorhabditis elegans*, *Dictyostelium discoideum*, *Naegleria fowleri*, *Tetrahymena thermophila*, and *Homo sapiens* representing mammals (Figure 3G)^27, 28, 29^. The alignment reveals that residues H3A1 – H3K18, including the H3K14 methylation site, are either invariant or highly conserved among host species. In contrast, the residues following H3K18 have weaker sequence homology, with amino acid insertions or deletions in certain species. Notably, residues H3A1 – H3K18 are specifically recognized by the ankyrin and SET domains of LegAS4 (Figure 3A), indicating that the LpHKMTs of *L. pneumophila* have evolved to methylate H3K14 by binding to a region of the histone H3 N-terminal tail that is highly conserved across diverse hosts.

### H3K14 site specificity of the SET domain

The structure of the LegAS4/AdoHcy/H3_[1-21]_ peptide ternary complex offers molecular insights into the H3K14 specificity of the LpHKMTs. Histone H3 residues K9 – R17 flanking the K14 site are bound through an extensive network of van der Waals interactions and hydrogen bonds with the substrate binding cleft of the enzyme’s SET domain (Figure 4A). The majority of the hydrogen bonds between histone H3 and the enzyme involve backbone interactions, with the remainder mediated by amino acid side chains. In the enzyme’s conformationally flexible switch region, the carboxamide group of Asn221 forms bidentate hydrogen bonds with the carbonyl oxygen of H3A15 and the guanidinium group of H3R17, while the hydroxyl group of Thr222 hydrogen bonds with the carbonyl group of H3G13. These interactions are enabled by conformational changes in the switch region induced by the enzyme binding to the histone H3 N-terminal tail (Figure 3D). In addition, the ε-amino group of Lys157 in LegAS4 engages in a hydrogen bond with the carbonyl oxygen of H3T11 (Figure 4A). Alanine substitutions of Lys157 and Thr222 in RomA yielded minor to modest alterations in nucleosome methylation relative to the WT enzyme (Figure 4B, S6A). In contrast, the RomA Asn221 to alanine mutant exhibited a notable decrease in activity. This mutation abolishes the bidentate hydrogen bonding to H3A15 and H3R17, potentially disrupting the conformational changes in the SET domain C-terminal loop that promote H3 binding (Figure 3D, 4A).

**Figure 4:**
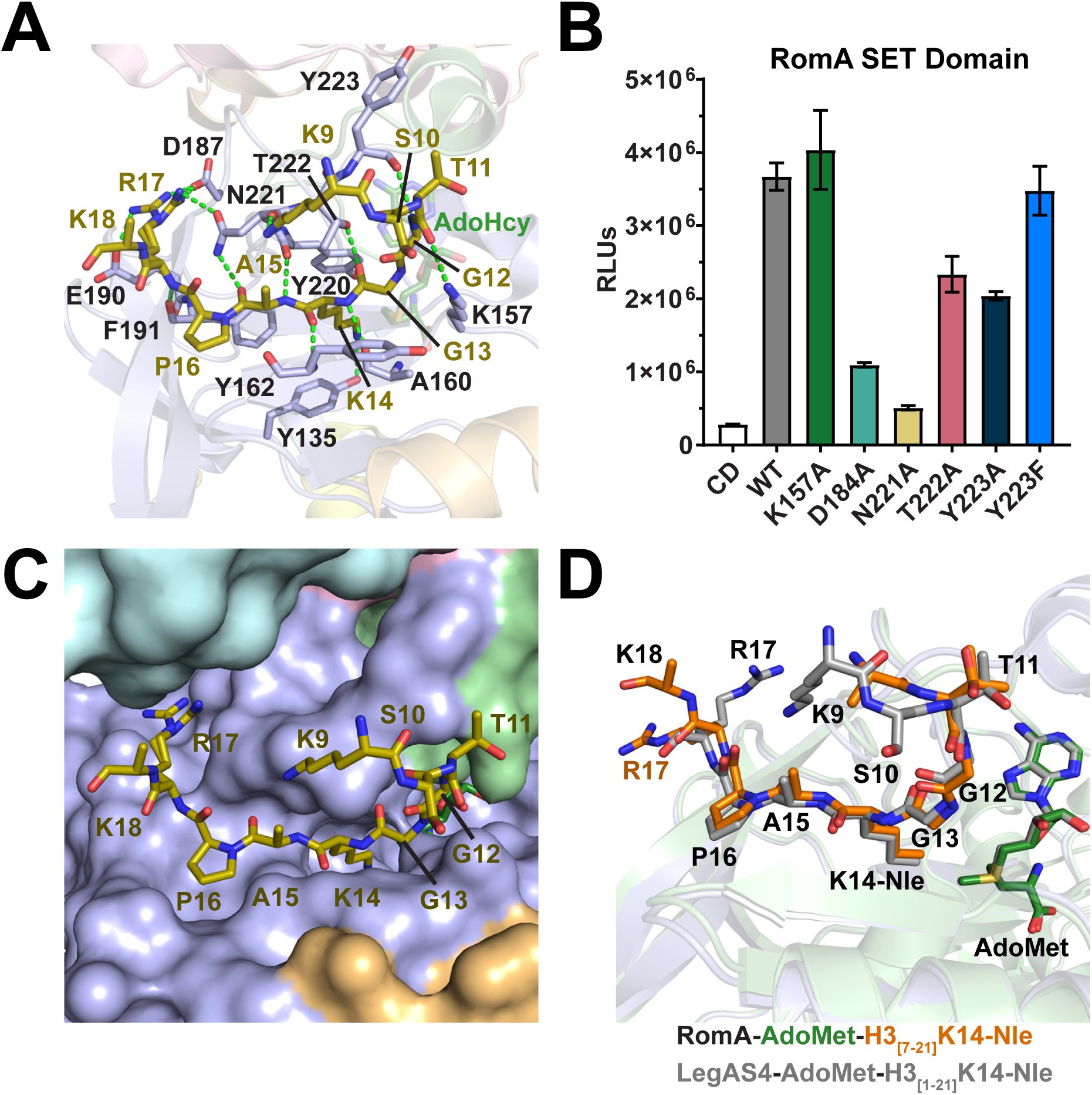
H3K14 Recognition by the SET domain. **A.** LegAS4 SET domain bound to AdoHcy (green carbon atoms) and H3K9 – H3K18 in the histone H3_[1-21]_ peptide (gold carbon atoms), with domains and subdomains colored as in Figure 3A. Hydrogen bonds are depicted as green dashes (Table S5). H3R17 adopts two alternative conformations. **B.** Luminescent methyltransferase assays with WT RomA, the RomA CD mutant, and RomA SET domain mutants using *X. laevis* ([H3.1]_2_) as substrates. Assays were performed in triplicate, and error bars represent standard deviation of the mean. **C.** Molecular surface of the SET domain and neighboring regions in LegAS4 bound to residues H3K9 – H3K18 in the histone H3_[1-21]_ peptide. **D.** Structural alignment of the SET domains in the RomA/AdoMet/H3_[7-21]_K14-Nle peptide (colored as in Figure 3E) and LegAS4/AdoMet/H3_[1-21]_K14-Nle peptide (gray) complexes.

Unique surface features in their SET domains enable the LpHKMTs to specifically recognize H3K14 and its flanking residues. H3G12 and H3G13 bind in a narrow groove on the enzymes’ surface (Figure 4C), with the contour of the groove inducing H3G12 and H3G13 to form a sharp kink by adopting dihedral angles that are generally disfavored for non-glycine amino acids. Further, the groove’s dimensions cannot accommodate large residues without inducing steric clashes, explaining the enzymes’ selectivity for H3G12 and H3G13. H3P16 fits snugly in a complementary-shaped pocket on the surface of the substrate binding cleft through van der Waals interactions. Superimposition of the SET domains from the RomA/AdoMet/H3_[7-21]_K14-Nle peptide and LegAS4/AdoHcy/H3_[1-21]_K14-Nle peptide complexes illustrates structurally homologous conformations of the histone H3 peptides, particularly residues H3G12 – H3R17, with the exception of the intrinsically flexible H3R17 side chain that adopts alternate conformations with the SET domains (Figure 4D, S4C). The conserved recognition mode observed for H3G12 – H3P16 in the LpHKMT structures concurs with an analysis of RomA lysine site specificity indicating the enzyme recognizes a G-<u>K</u>-X-P/A sequence motif (where <u>K</u> is the methylation site and X is any amino acid)^30^. Notably, this sequence motif is unique to H3K14 relative to other potential lysine methylation sites in the core histone N-terminal tails, illuminating LpHKMT site specificity (Figure S6B).

### Recognition of histone H3 N-terminus by the LpHKMTs

The structure of LegAS4 bound to the histone H3_[1-21]_ peptide reveals that histone residues H3A1 – H3R8 engage in a novel set of interactions with the ankyrin domain and the interface between the SET and ankyrin domains (Figure 5A). Notably, residues H3T3 – H3A7 adopt a helical conformation stabilized by a network of intramolecular hydrogen bonds within the H3_[1-21]_ peptide backbone and the side chains of H3T3 and H3T6, in addition to intermolecular hydrogen bonds and van der Waals interactions with residues in the ankyrin domain. Due to this conformation, H3A1, H3R2, H3K4, and H3Q5 engage in a network of hydrogen bonds with residues in the ankyrin and SET domains (Figure 5A-C). H3A1 is bound within a pocket at the interface of the ankyrin and SET domains in which its α-ammonium cation forms a salt bridge interaction with the carboxylate side chain of Asp184 in the SET domain. The H3R2 side chain binds in a surface cleft where its guanidinium group forms a hydrogen bond with the side chain of Tyr223 from the flexible loop of the SET domain and a salt bridge with Asp438 in the ankyrin repeats. Notably, upon H3 binding, Tyr223 undergoes a large conformational change that orients its side chain in the surface cleft for interactions with H3R2 (Figure 3D and Movie S1). The side chain of H3K4 inserts into a deep channel at the interface of the last ankyrin repeat and C-cap region, whose hydrophobic residues engage in van der Waals interactions with the aliphatic side chain of H3K4 (Figure 5C). At the base of the channel, the H3K4 ε-amino group forms a salt bridge with the carboxylate side chain of Asp440 and hydrogen bonds with the carbonyl oxygen of Phe431 and the side chain of Asn480, anchoring the H3K4 side chain within the lysine binding channel. Finally, H3Q5 engages in a hydrogen bond with the side chain of Thr432 on the surface of the ankyrin repeats.

**Figure 5:**
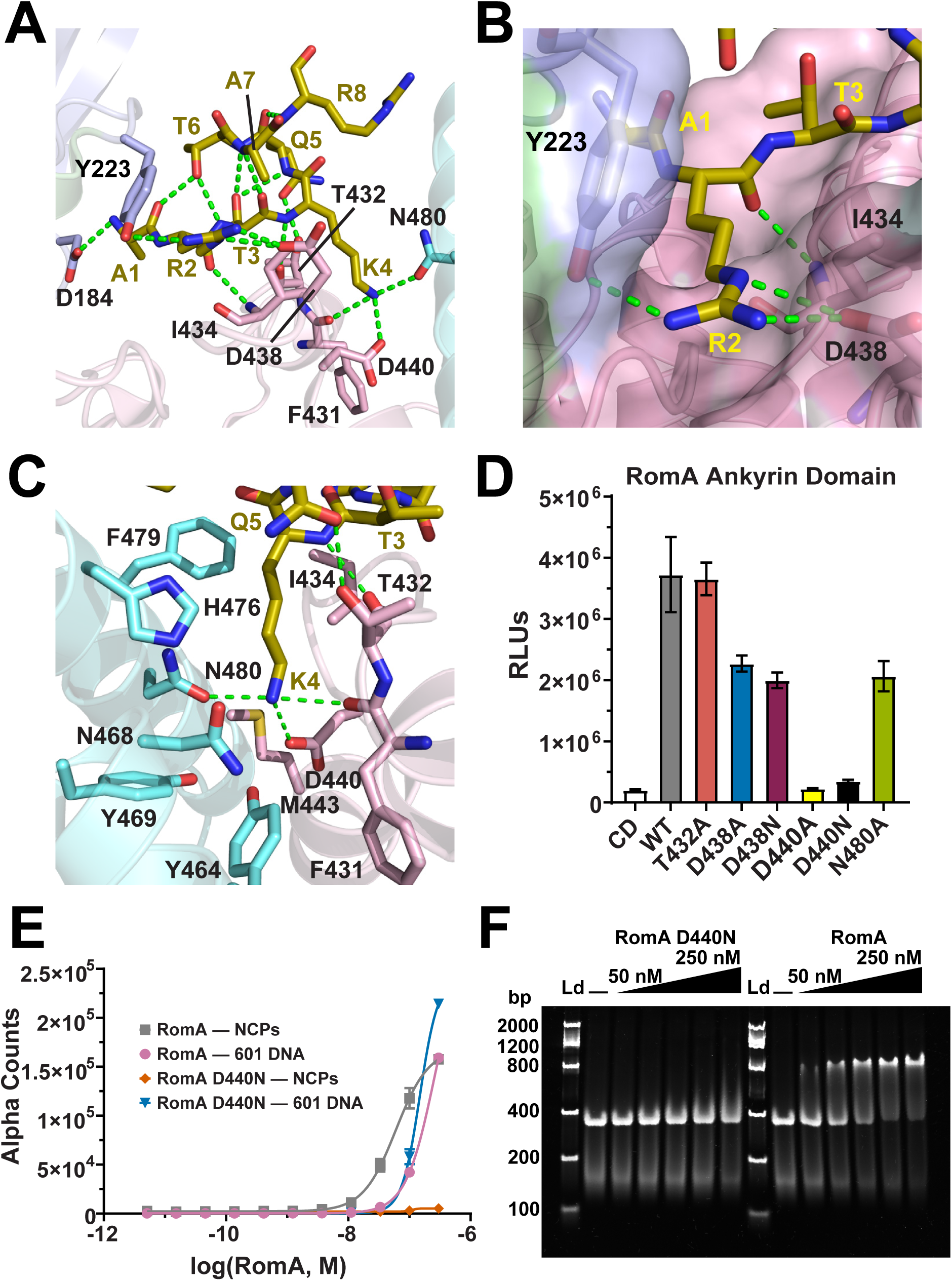
Recognition of histone H3 N-terminal tail by the Ankyrin repeat domain. **A.** Hydrogen bonding between the LegAS4 Ankyrin repeat domain and histone H3 residues A1 – R8 (Table S5). **B.** Histone H3R2 binds within a surface cleft through hydrogen bonding with residues in the SET and Ankyrin repeats domains. **C.** Histone H3K4 is inserted in a deep pocket in the LegAS4 Ankyrin domain. The H3K4 side chain is stabilized through hydrogen bonding with three residues in the base of the pocket. **D.** Luminescent methyltransferase assays with WT RomA, RomA CD mutant, and RomA Ankyrin domain mutants using *X. laevis* ([H3.1]_2_) NCPs. **E.** Captify-Alpha assays for WT RomA and D440N mutant binding to human ([H3.1]_2_) and 147 bp 601 DNA. **F.** EMSAs of WT RomA and the D440N mutant with recombinant *X. laevis* ([H3.1]_2_).

The observations that H3A1, H3R2, H3K4, and H3Q5 directly interact with LegAS4 spurred us to examine the roles of these residues in histone H3 engagement. In the H3A1 binding pocket of RomA, mutation of Asp184 to alanine (D184A) impaired nucleosome methylation (Figures 4B, 5A, S6A), demonstrating the importance of the salt bridge between Asp184 and H3A1. Conservative and alanine mutations of Tyr223 (Y223F and Y223A) and Asp438 (D438N and D438A) in RomA that disrupt hydrogen bonding to the H3R2 guanidinium group diminished methyltransferase activity to varying degrees, indicating that H3R2 interactions within its surface cleft on the ankyrin repeats also facilitate substrate engagement (Figures 4B, 5B, 5D, S6A, S7A). In comparison to H3A1 and H3R2, substitutions of residues that hydrogen bond to the ε-amino group of H3K4 had a more pronounced effect on RomA activity. Mutation of Asn480 to alanine (N480A) in RomA reduced nucleosome methylation approximately two-fold, whereas substitutions of Asp440 by alanine (D440A) and asparagine (D440N) abolished activity (Figure 5C, 5D, S7A). Captify-Alpha assays and EMSAs revealed that the RomA D440N mutation abrogated nucleosome binding, confirming that the salt bridge between Asp440 and the H3K4 side chain is crucial for histone H3 recognition (Figure 5E, 5F, S7B). Conversely, the association of RomA D440N with the 147 bp 601 DNA was unaffected, consistent with the interactions between the enzyme’s NLS and the DNA being unaffected by this mutation. Finally, we examined the hydrogen bonding between H3Q5 and Thr432 in RomA and observed that an alanine substitution of this residue (T432A) had no appreciable impact on nucleosome methylation, suggesting that this interaction is dispensable (Figure 5C, 5D, S7A). Collectively, these findings illustrate that interactions between the LpHKMTs and N-terminal residues of histone H3 are pivotal to nucleosome engagement and H3K14 methylation, most particularly the binding of H3K4 within the lysine binding channel of the ankyrin domain.

### The Ankyrin domain reads the modification states of H3R2 and H3K4

Our findings that various H3R2 and H3K4 modifications alter nucleosome engagement by RomA prompted us to investigate the mechanism by which the enzyme differentiates these PTMs (Figure 2D). Expanding on the earlier panel profiling, we performed Captify-Alpha assays titrating RomA against various H3R2-modified nucleosomes and observed comparable affinity toward ([H3R2un]_2_) and ([H3R2me1]_2_) and diminished binding to ([H3R2me2a]_2_) and ([H3R2me2s]_2_) (Figure 6A). Likewise, RomA and LegAS4 methylated ([H3R2un]_2_) and ([H3R2me1]_2_) with similar efficiency, while methylation of ([H3R2me2a]_2_) and ([H3R2me2s]_2_) was reduced approximately twofold (Figure 6B, 6C, S8A, S8B). Crystal structures of LegAS4 in complex with histone H3_[1-21]_ peptides possessing different H3R2 methylation states illustrate that the side chains of H3R2me1 and H3R2me2s adopt conformations analogous to H3R2un and engage in similar patterns of interactions with the side chains of Tyr223 and Asp438 in LegAS4 (Figure 5B, 7A – 7D and Table S3, S4). Conversely, the guanidinium moiety of H3R2me2a is flipped relative to the orientations of the H3R2 side chains in the other structures, resulting in a loss of hydrogen bonding to Tyr223 while maintaining a salt bridge with the carboxylate group of Asp438 (Figure 7E, 7F). Despite their alternative conformations, the H3R2me1, H3R2me2a, and H3R2me2s share common binding modes in that their respective methyl groups generally project outward from the H3R2 binding cleft towards solvent, thereby mitigating the loss of hydrogen bonding and other interactions with residues in this surface cleft (Figure 7). To explore a more dramatic change than arginine methylation, we truncated the first two residues in histone H3 N-terminus (H3βN2) or mutated H3R2 to alanine (H3R2A). As anticipated, each mutant disrupted nucleosome binding and methylation by the LpHKMTs, indicating that H3R2 recognition is crucial to H3K14 methylation by the enzymes (Figure 6, S8C).

**Figure 6:**
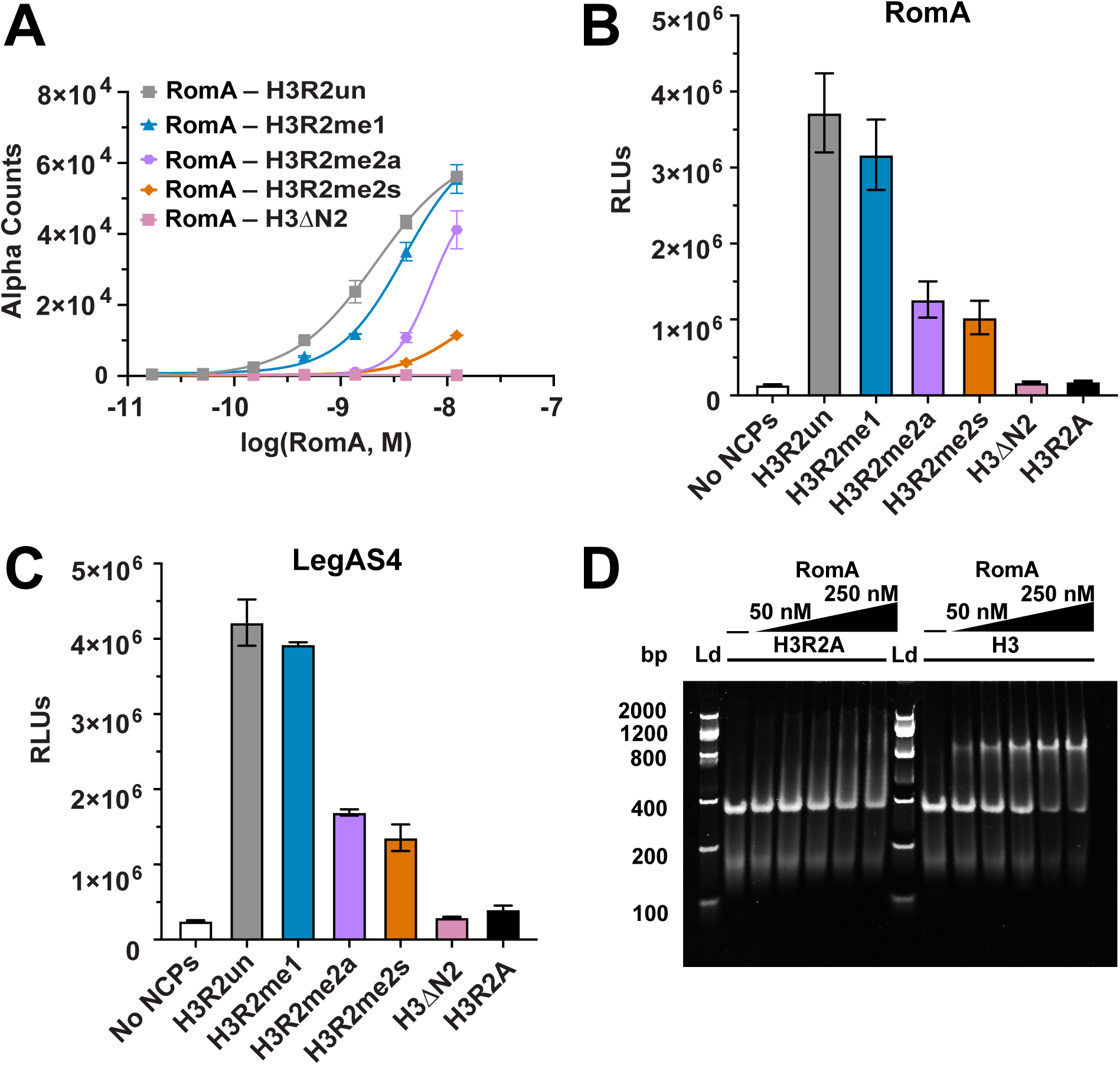
Recognition of H3R2 methylation by the LpHKMTs. **A.** Captify-Alpha assays of RomA with human ([H3R2un]_2_), ([H3R2me1]_2_), ([H3R2me2a]_2_), ([H3R2me2s]_2_), or ([H3ΔN2]_2_) NCPs. **B, C.** Luminescent methyltransferase assays with RomA and LegAS4 with ([H3R2un]_2_), ([H3R2me1]_2_), ([H3R2me2a]_2_), ([H3R2me2s]_2_), ([H3ΔN2]_2_), or ([H3.1R2A]_2_) (Table S1). **D.** EMSAs analyzing the binding of RomA to *X. laevis* ([H3.1]_2_) and ([H3.1R2A]_2_) NCPs.

**Figure 7:**
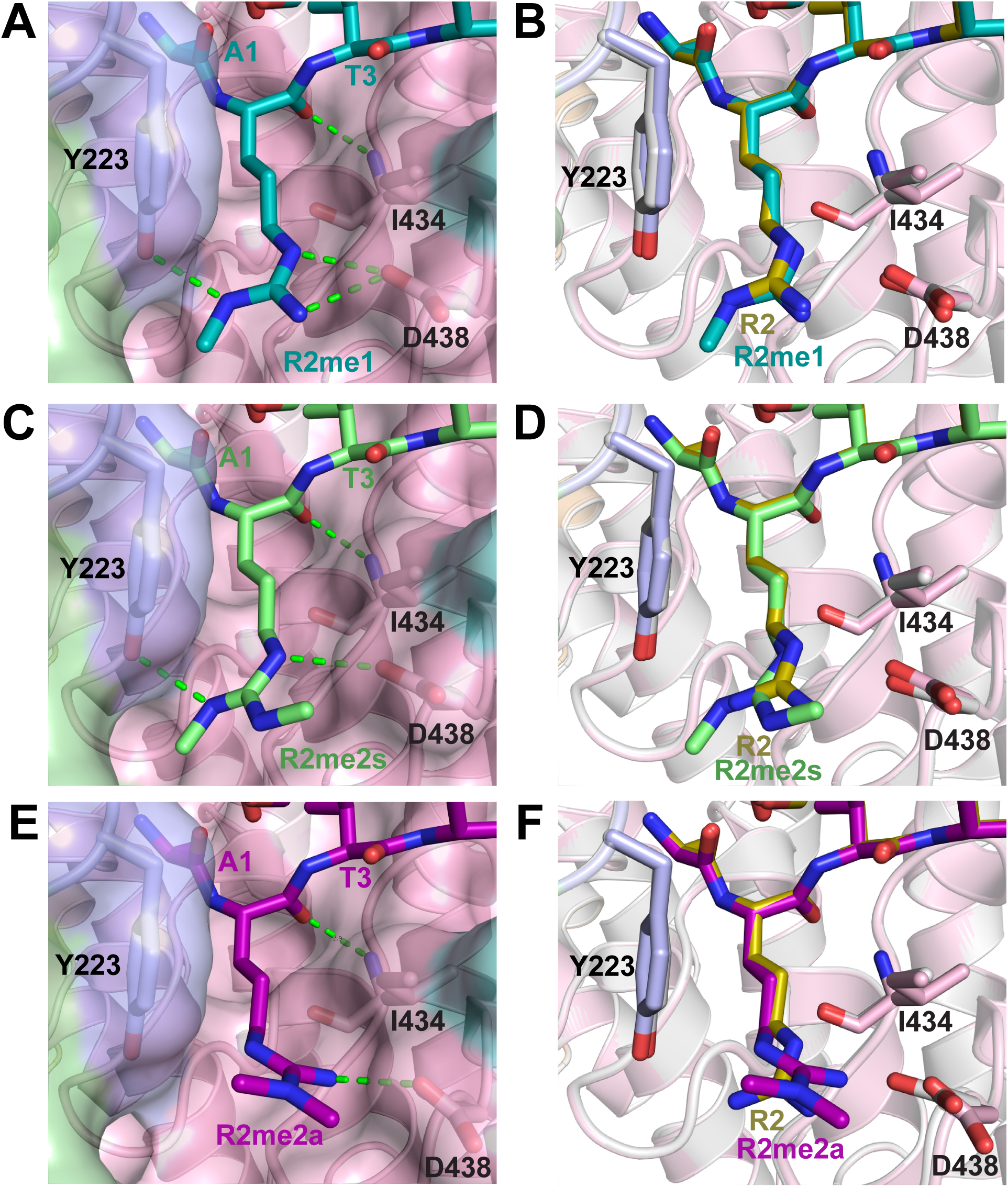
Molecular basis for the recognition of the H3R2 methylated states by LegAS4. **A.** H3R2me1 (teal carbon atoms) bound within the H3R2 binding cleft of LegAS4. Hydrogen bonding between H3R2me1 and residues in LegAS4 are depicted as green dashes. **B.** Superimposition of the structures of LegAS4 complexes with the histone H3_[1-21]_ peptide (light gray cartoon) and H3_[1-21]_R2me1 peptide (cartoon colored as Figure 3A) illustrating the overlaid binding modes of H3R2 (gold carbon atoms) and H3R2me1. **C,D.** Structure of H3R2me2s (light green carbon atoms) bound in the H3R2 binding cleft, as depicted in panels **A** and **B**, respectively. **E,F.** H3R2me2a (violet carbon atoms) bound within the H3R2 binding cleft illustrated as in panels **A** and **B**, respectively.

Variations in the dimensions, solvent exposure, and interactions within the H3R2 surface cleft and H3K4 binding channel define the propensity of LpHKMTs to discriminate the PTM landscape at these positions. Here H3K4 PTMs often exhibited a more pronounced effect on nucleosome engagement and H3K14 methylation. RomA bound ([H3K4un]_2_) and ([H3K4me1]_2_) with similar affinities but displayed minimal association with ([H3K4me2]_2_), ([H3K4me3]_2_), or ([H3K4ac]_2_) (Figure 8A). Consistent with these results, RomA and LegAS4 efficiently methylated ([H3K4un]_2_) and ([H3K4me1]_2_) but not ([H3K4me2]_2_), ([H3K4me3]_2_), or ([H3K4ac]_2_) (Figure 8B, S9A – S9C). Further confirming the importance of the H3K4 side chain to the LpHKMTs, RomA and LegAS4 methylation of ([H3.1K4A]_2_) was reduced to near background levels (Figure 8B, S9A – S9C), and orthogonal EMSAs confirmed the diminished binding of RomA to these nucleosomes (Figure 8C, S9D). Lastly, we observed no nucleosome engagement by RomA D440N in Captify-Alpha assays (Figure S9E), consistent with the importance of this residue in H3K4 recognition. Collectively, these results demonstrate that LpHKMT reading of H3K4 and its modification state is crucial for nucleosome engagement and H3K14 methylation.

**Figure 8:**
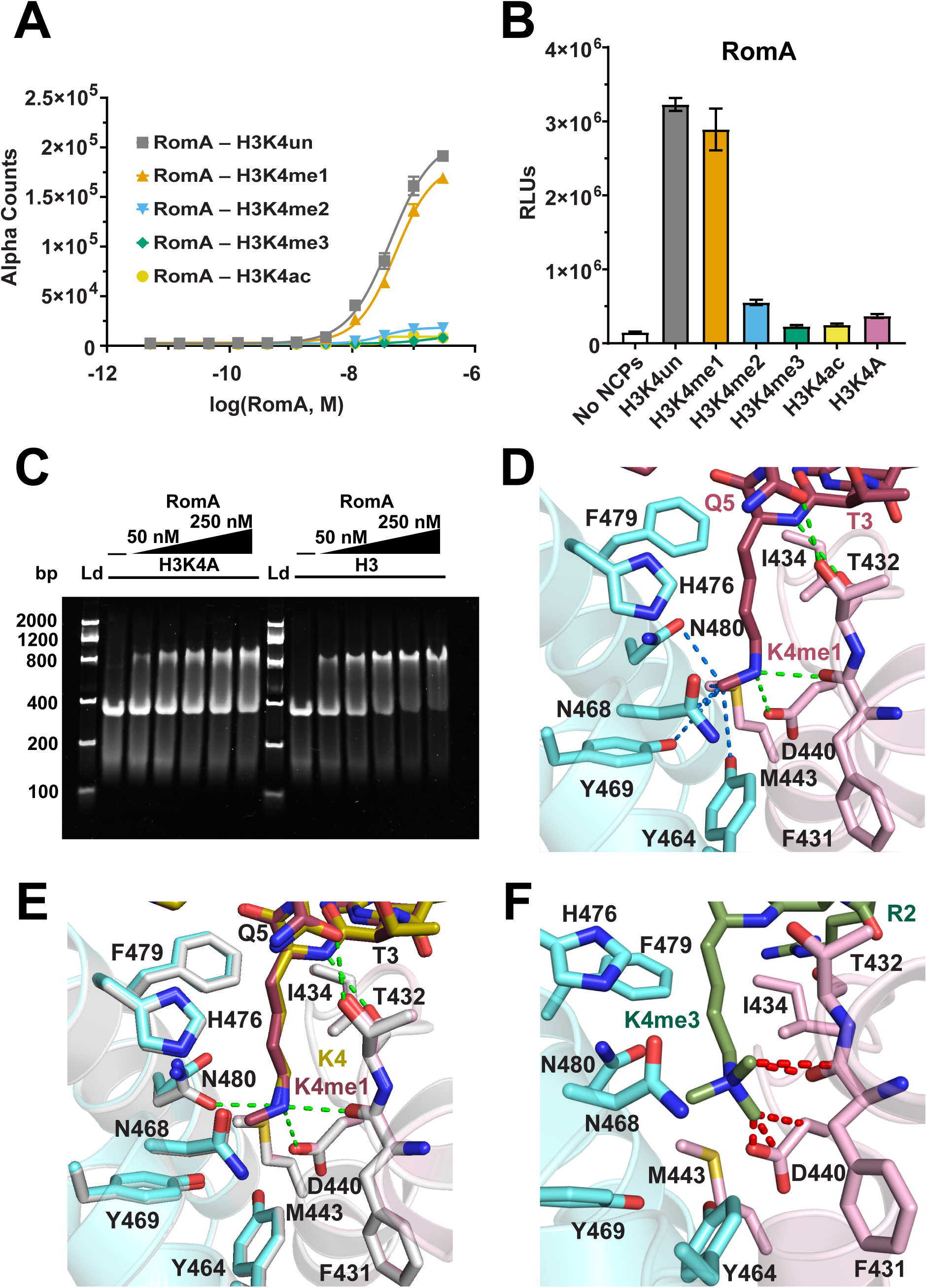
Recognition of the H3K4 modification state by the Ankyrin repeat domain. **A.** Captify-Alpha assays measuring the association of RomA with H3K4-modified NCPs (Table S1). **B.** Luminescent methyltransferase assays of RomA with ([H3K4un]_2_), ([H3K4me1]_2_), ([H3K4me2]_2_), ([H3K4me3]_2_), ([H3K4ac]_2_), or ([H3.1K4A]_2_) NCPs. **C.** EMSAs of RomA with *X. laevis* ([H3.1]_2_) and ([H3.1K4A]_2_) NCPs. **D.** Insertion of the H3K4me1 side chain (raspberry carbon atoms) into the H3K4 binding pocket of LegAS4 with hydrogen bonds to the lysine ε-amino group (green dashes) and van der Waals contacts with the methyl group (blue dashes) illustrated. **E.** Overlay of the H3K4 binding pockets of the LegAS4/H3_[1-21]_K4un peptide (gray with gold carbons for H3K4) and LegAS4/H3_[1-21]_K4me1 peptide (depicted as in **D**) complexes. **F.** Model of H3K4me3 within the H3K4 binding pocket of LegAS4 with steric clashes denoted as red dashes.

To elucidate the molecular basis by which the LpHKMTs recognize different H3K4 PTMs, we determined the crystal structure of LegAS4 bound to AdoHcy and a histone H3_[1-21]_K4me1 peptide at 1.75 Å resolution (Figure 8D and Table S3, S4). The structure reveals that the H3K4me1 side chain is bound within the H3K4 binding channel through hydrogen bonding and van der Waals interactions similar to the binding mode of H3K4, with the exception that the hydrogen bond between the H3K4 ε-amino group and Asn480 side chain in LegAS4 is abrogated by the methyl group. In a shallow cavity at the base of H3K4 binding pocket, the methyl group is bound via van der Waals interactions with Met 443, Tyr464, Asn468, Tyr469, and Asn480 in LegAS4. Superimposing the structures of LegAS4 bound to H3K4 and H3K4me1 illustrates that the enzyme’s residues composing the H3K4 binding channel adopt essentially identical conformations, with the exception of the carboxamide side chain of Asn480 that pivots to form a van der Waals interaction with the methyl group of H3K4me1 (Figure 8E).

We next examined the mechanism by which the LpHKMTs distinguish H3K4me1 from bulkier H3K4 PTMs. Modeling of H3K4me3 based on the coordinates of the H3K4me1 side chain reveals that the two additional methyl groups in the trimethyllysine sterically clash with the carboxylate group of Asp440 and the carbonyl oxygen of Phe431 (Figure 8F). Notably, these two residues hydrogen bond with the ε-amino group of H3K4 and H3K4me1; thus, steric occlusion may serve to discriminate against the binding of H3K4me2, H3K4me3, and H3K4ac. Taken together, the hydrogen bonding network and dimensions of the H3K4 binding channel in the ankyrin domain confer selectivity in recognizing H3K4 and H3K4me1 but impede the binding of bulkier lysine PTMs by steric hindrance.

## Discussion

These studies furnish new insights into nucleosome engagement and histone H3 recognition by the LpHKMT enzymes. Nucleosome engagement is driven by the enzymes’ binding to the first 18 residues in the N-terminal tail of histone H3 and electrostatic interactions between their lysine-rich NLS motifs and the nucleosomal DNA. Basic regions on the surfaces of other HKMTs or HKMT complexes, including DOT1L, MLL/SET1, NSD, PRC2, and SETD2, engage in electrostatic interactions with nucleosomal DNA, illustrating a common mechanism for chromatin _association_^31, 32, 33, 34, 35, 36, 37, 38, 39, 40, 41, 42, 43, 44, 45, 46, 47^. This engagement mechanism concurs with the ability of RomA to bind nucleosomes harboring diverse histone variants or PTMs with equivalent affinity and thus write H3K14 methylation across the host cell genome during *L. pneumophila* infection^6, 13, 14^.

The crystal structures and biochemical characterization of LegAS4 and RomA reveal a novel mechanism by which the LpHKMTs read and write PTMs on histone H3. The enzymes’ unique V-shaped architecture promotes bipartite binding of the histone H3 N-terminal tail, enabling simultaneous engagement of H3A1 – H3R8 by the ankyrin domain and H3K9 – H3K18 by the SET domain. Histone H3 binding to the LpHKMTs induces a conformational change in the substrate binding cleft of their SET domains, facilitating recognition of the H3K14 methylation site. In contrast, the ankyrin domain does not undergo large scale conformational changes during histone H3 binding. Instead, residues H3T3 – H3R8 adopt a helical conformation that interacts with the ankyrin repeats and the interface between the ankyrin and SET domains.

A structural survey of histone readers reveals that this helical conformation is not unique to the bacterial LpHKMTs, as certain eukaryotic plant homeodomains (PHDs) induce similar helicity in the N-terminus of histone H3. These readers include the PHD domain of BAZ2A in the NoRC chromatin remodeling complex, the tandem Tudor and PHD domains of the UHRF1 E3 ubiquitin ligase, the double PHD finger (DPF) of the MOZ histone acetyltransferase, and the DPF of the BAF45C subunit of the BAF chromatin remodeler^48, 49, 50, 51, 52^. Notably, the structures of these reader domains illustrate that they engage in hydrogen bonding to the α-amino group of H3A1 and the side chains of H3R2 and H3K4, analogous to the recognition of these residues by LegAS4.

To our knowledge, the ankyrin repeat domain of the LpHKMTs represents the first example of a bacterial histone reader for eukaryotic histone PTMs. Studies of other histone methyltransferases expressed by bacterial pathogens, including *Bacillus anthracis*, *Burkholderia thailandensis*, *Chlamydia trachomatis*, and *Mycobacterium tuberculosis,* have reported their respective substrate selectivities but have not revealed histone reader activities^7, 53, 54, 55, 56^. Ankyrin repeats represent a ∼33 residue helical motif that frequently mediates protein-protein interactions. The only described example as a histone reader is in the metazoan family of H3K9-specific euchromatic histone methyltransferases (EHMTs) that include mammalian G9A and its homolog G9A-like protein (GLP)^57, 58^. Histone PTM reading by the LpHKMTs and EHMTs differs in several respects. Notably, the ankyrin repeat domain in the EHMTs possesses a canonical aromatic cage that selectively recognizes H3K9me1/2 through cation-π interactions and an electrostatic interaction with a glutamate residue^59^. In the LpHKMTs, the H3K4 binding channel in the ankyrin domain selectively binds H3K4un and H3K4me1 through predominantly van der Waals contacts and hydrogen bonding with the lysine side chain and discriminates against H3K4me2/3 through steric occlusion. Further, the ankyrin domain is distal to the SET domain in the EHMTs^59^ but adjacent in the LpHKMTs. These differences in reader and writer domain arrangement result in distinct modes of chromatin engagement. The distal arrangement of the ankyrin and SET domains in EHMT heterodimers respectively enables *trans* recognition of H3K9me1/2 and H3K9 in different histone H3 N-terminal tails in the same nucleosome or neighboring nucleosomes^60^. Conversely, the ankyrin and SET domain architecture in the LpHKMTs respectively promotes *cis* recognition of H3K4un/me1 and H3K14 in the same histone H3 tail. This binding mode is crucial to the function of the LpHKMTs, as ankyrin domain mutations that disrupt H3K4 binding also abrogate H3K14 methylation.

The discovery that the ankyrin domain specifically recognizes H3K4un and H3K4me1 before methylating H3K14 has important ramifications for localizing the LpHKMTs in chromatin. Notably, enhancer elements of genes are frequently marked with H3K4me1, whereas transcriptionally active loci, including promoter regions, are enriched in H3K4me3^61^. Prior chromatin immunoprecipitation sequencing (ChIP-Seq) of *L. pneumophila*-infected host cells indicated an enrichment of H3K14 methylation at the transcription start sites (TSSs)^6^ that are marked by H3K4me3 at actively transcribed genes. Such a distribution would appear to directly conflict with our structural and biochemical data demonstrating that RomA and LegAS4 are unable to bind and methylate H3K4me3 nucleosomes. We note that LpHKMT-catalyzed H3K14 methylation at H3K4me1-enriched transcriptional enhancers has the potential to antagonize H3K14ac, a PTM associated with chromatin remodeling and transcriptional activation^62, 63^. Future studies will be required to resolve the host loci that undergo H3K14 methylation during *L. pneumophila* infection and how this modification mechanistically contributes to pathogenesis.

RomA and LegAS4 are key effectors in *L. pneumophila* infection, where deletion of their cognate genes impairs intracellular replication in both protozoan and metazoan host cells^6^. This phenotype contrasts with the genetic knockout of numerous *L. pneumophila* effector genes that yield little to no effect on intracellular replication in host cells presumably due to functional redundancy among effectors^64^. The intrinsic ability of the LpHKMTs to recognize a highly conserved sequence motif in the histone H3 N-terminal tail – reading nucleosomal H3K4 and methylating H3K14 – provides a molecular explanation for their conserved function in diverse protozoan and metazoan hosts. These findings lay the groundwork for further exploring chromatin engagement and modification by the LpHKMTs and other bacterial effector proteins.

## Acknowledgements

The authors thank Chimi Sherpa for assistance with nucleosome reconstitution, Dr. Joel Swanson for insights in culturing BMDMs, Dr. Uhn-Soo Cho for advice on nucleosome preparation, and Dr. Robert Fick for assistance in rendering structural figures. In addition, we posthumously acknowledge Dr. William L. Smith for his support and Dr. Henriette Remmer for assistance in ordering peptides. We also wish to thank staff at the Life Sciences Collaborative Access Team (LS-CAT) at the Advanced Photon Source Synchrotron for support with X-ray data collection. This work is supported by National Institutes of Health grant R01AI160067 and intramural funding from the University of Michigan Medical School Biomedical Research Council (to R.C.T.). EpiCypher is supported by NIH grants R44GM117683, R44CA214076, R44GM136172 and R44GM116584. S.G. is supported by a National Science Foundation Graduate Research Fellowship DGE 2241144. This research used resources of the Advanced Photon Source, a U.S. Department of Energy (DOE) Office of Science User Facility operated for the DOE Office of Science by Argonne National Laboratory under Contract No. DE-AC02-06CH11357. Use of the LS-CAT Sector 21 was supported by the Michigan Economic Development Corporation and the Michigan Technology Tri-Corridor (Grant 085P1000817).

## Materials and Methods

Please see the Supplementary Information.

