## Supplementary Information for "Molecular Basis of Histone H3 Reading and Writing by *Legionella pneumophila* SET Domain Lysine Methyltransferases"

- **SUPPLEMENTARY FIGURES**

**Figure S1:** LegAS4 and RomA sequence homology and substrate specificity.

**Figure S2:** LegAS4 methylates H3K14 in cells and *in vitro*.

**Figure S3:** Nucleosome recognition by RomA and LegAS4.

**Figure S4:** Binding of histone H3 peptides and AdoHcy to LegAS4.

**Figure S5:** ITC analysis of substrate binding to RomA.

**Figure S6:** H3K14 recognition by the SET domain.

**Figure S7:** Recognition of H3 N-terminal tail by the Ankyrin repeat domain.

**Figure S8:** Recognition of H3R2 methylation by the LpHKMTs.

**Figure S9:** Reading of H3K4 modifications by the Ankyrin repeat domain.

- **SUPPLEMENTARY METHODS**

- **SUPPLEMENTARY TABLES**

**Table S1:** Histone octamers and nucleosomes.

**Table S2:** LpHKMT constructs.

**Table S3:** Crystallographic and refinement data.

**Table S4:** Histone H3 peptides.

**Table S5:** LegAS4-histone H3 hydrogen bonding.

**Table S6:** Strains.

**Table S7:** Primers.

**Table S8:** Reagents.

**Table S9:** Plasmids.

**Table S10:** Antibodies.

### SUPPLEMENTARY FIGURES:

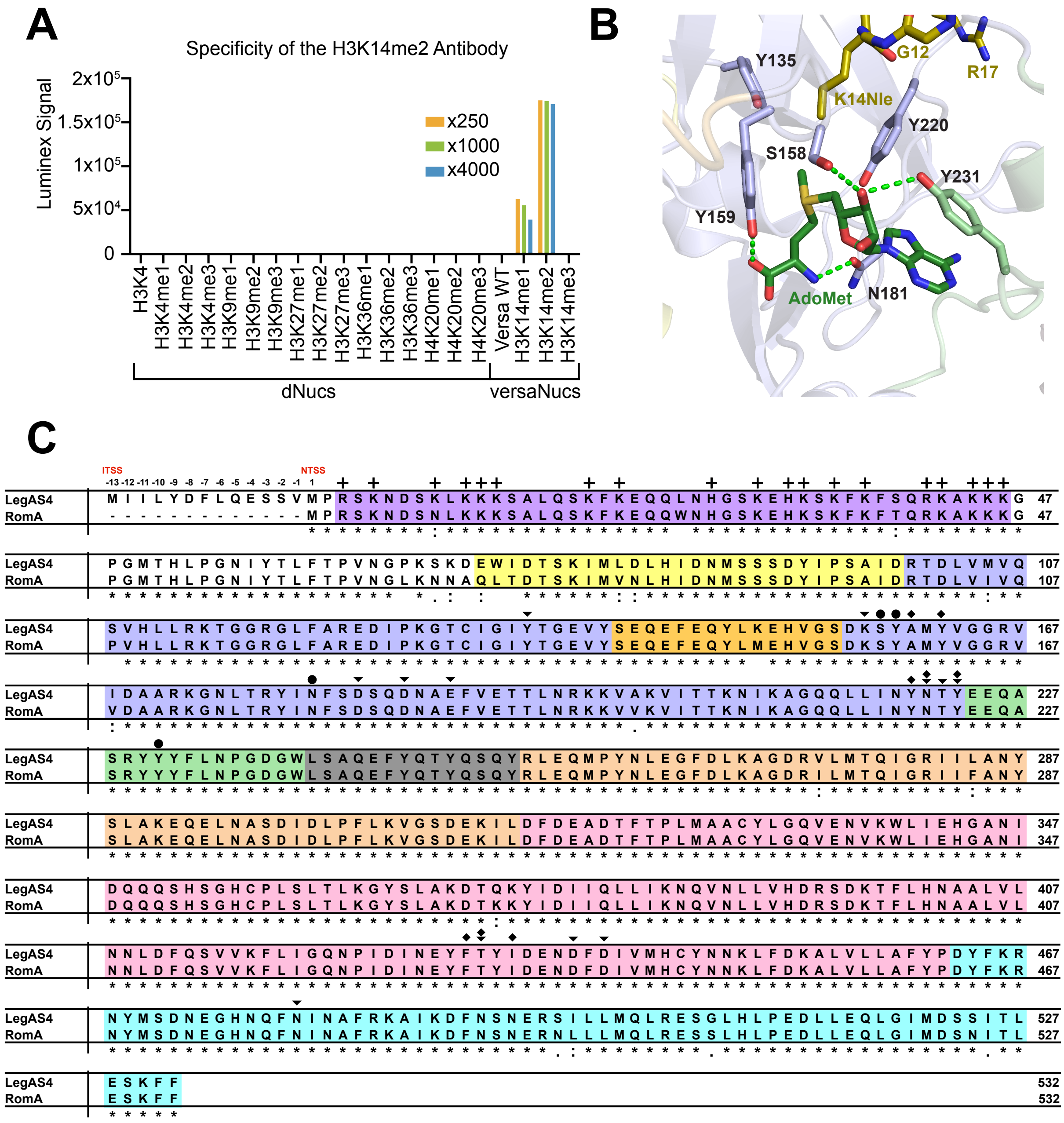

**Figure S1: LegAS4 and RomA sequence homology and substrate specificity. A.** Captify™-Luminex screening of anti-H3K14me2 (PIMA3059) against a panel of histone methyllysine-bearing nucleosome core particles (NCPs) illustrates its relative preference for H3K14 methylation. Inset numbers denote antibody dilution factor. NCPs utilized a combination of modified designer nucleosomes (dNucs™) and versatile nucleosome assembly (versaNuc™). **B.** The six residues (black labels) mutated to generate the hextuple catalytic defective (CD) mutant depicted by cartoon and stick representation of the active site of LegAS4. Hydrogen bonds are depicted as green dashes. **C.** Pairwise sequence alignment using Clustal Omega of the LpHKMT orthologs LegAS4 and RomA from the *L. pneumophila* Philadelphia-1 and Paris strains, respectively. The initial and new translation start sites (ITSS and NTSS) reported for LegAS4 are labeled at its N-terminus. Colors indicate the nuclear localization signal (NLS; purple), nSET (yellow), SET (violet), iSET (orange), cSET (light green), linker (grey), ankyrin N-cap (wheat), ankyrin repeats (light pink), and ankyrin C-cap (aquamarine) subdomains. Residues in the enzymes that engage in hydrogen bonding with histone H3 through their main and/or side chains are denoted as diamonds (♦) and downward triangles (▼), respectively. Circles (●) signify amino acids that form hydrogen bonds with the substrate S-adenosylmethionine (AdoMet) or the product S-adenosylhomocysteine (AdoHcy). Residues that are identical (*), conserved (:), or semi-conserved (.) between LegAS4 and RomA are denoted under the sequence alignment.

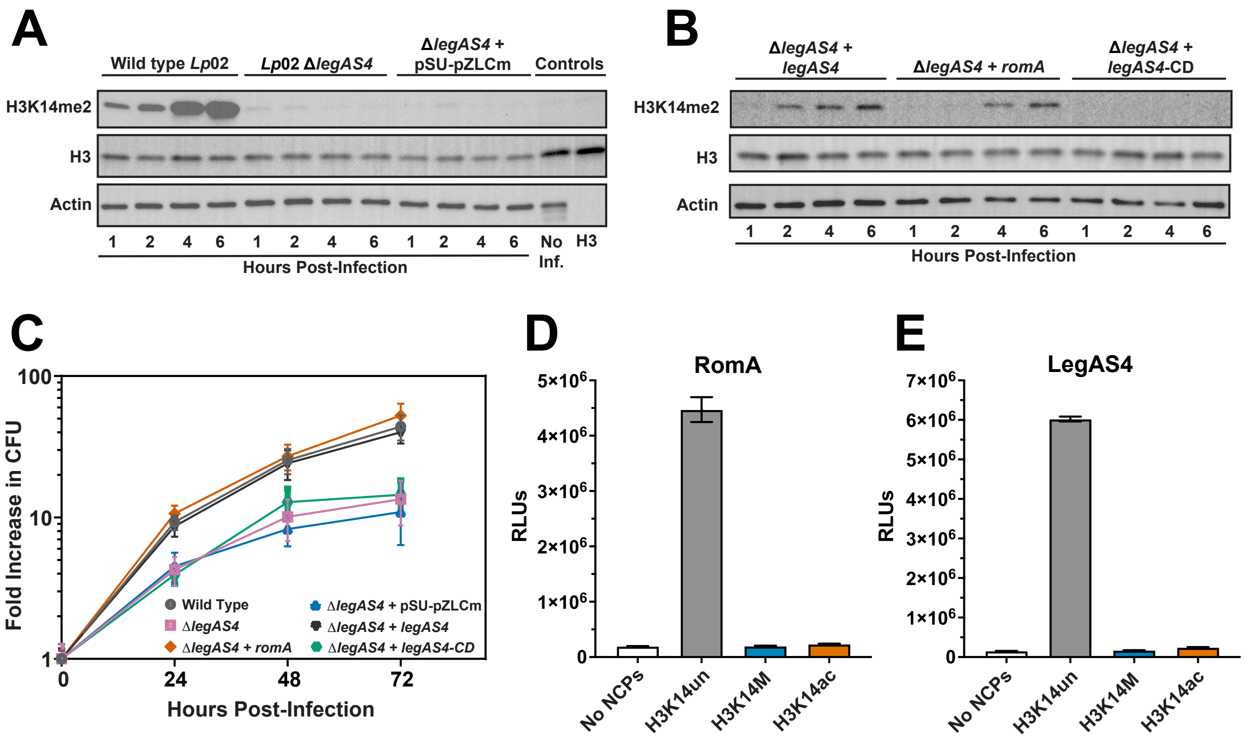

**Figure S2: LegAS4 methylates H3K14 in cells and *in vitro*. A, B, C.** Second biological replicates of Figure 1A, 1B, and 1C. **D, E.** Second replicates of the luminescent methyltransferase assays in Figures 1D and 1E.

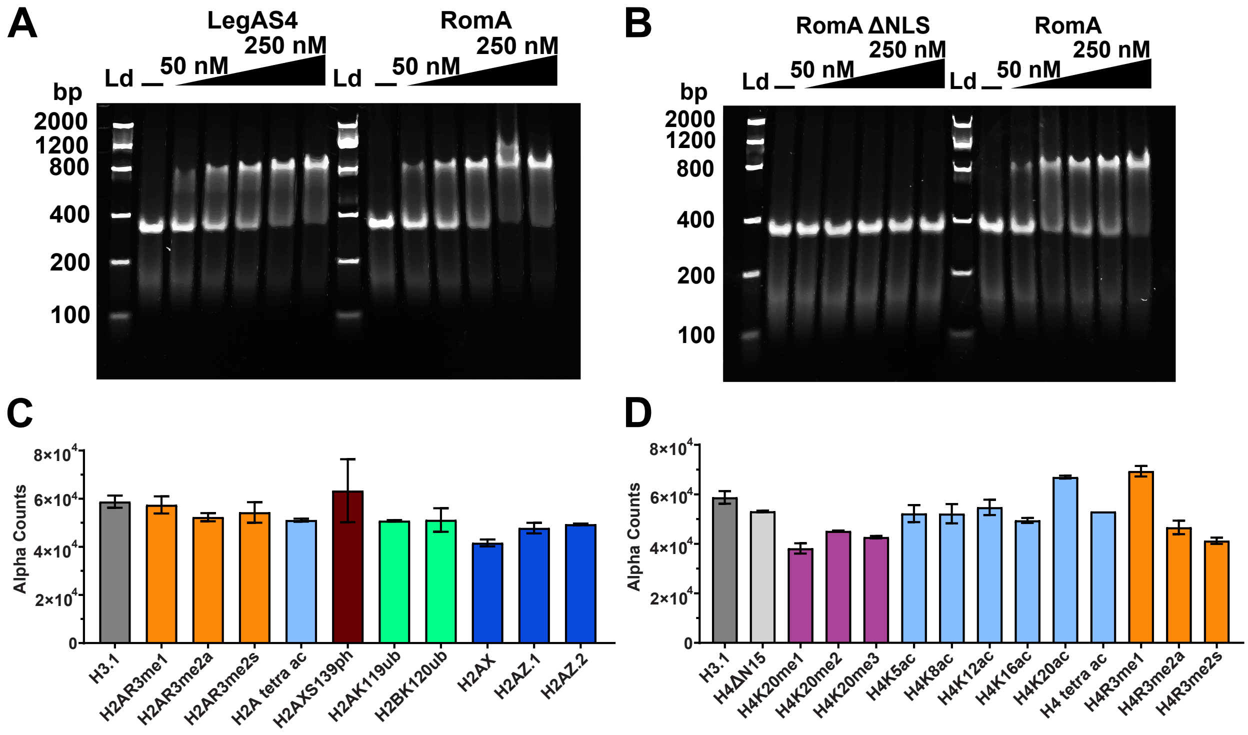

**Figure S3: Nucleosome recognition by RomA and LegAS4. A, B.** Second replicates of the EMSAs displayed in Figure 2B and 2C. **C, D.** Captify-Alpha assays screening the association of RomA with human NCPs bearing histone H2A or H2B PTMs, or H2A variants (**C**), or H4 PTMs (**D**). For comparison, NCPs possessing histone H3.1 are included as a positive control (Table S1).

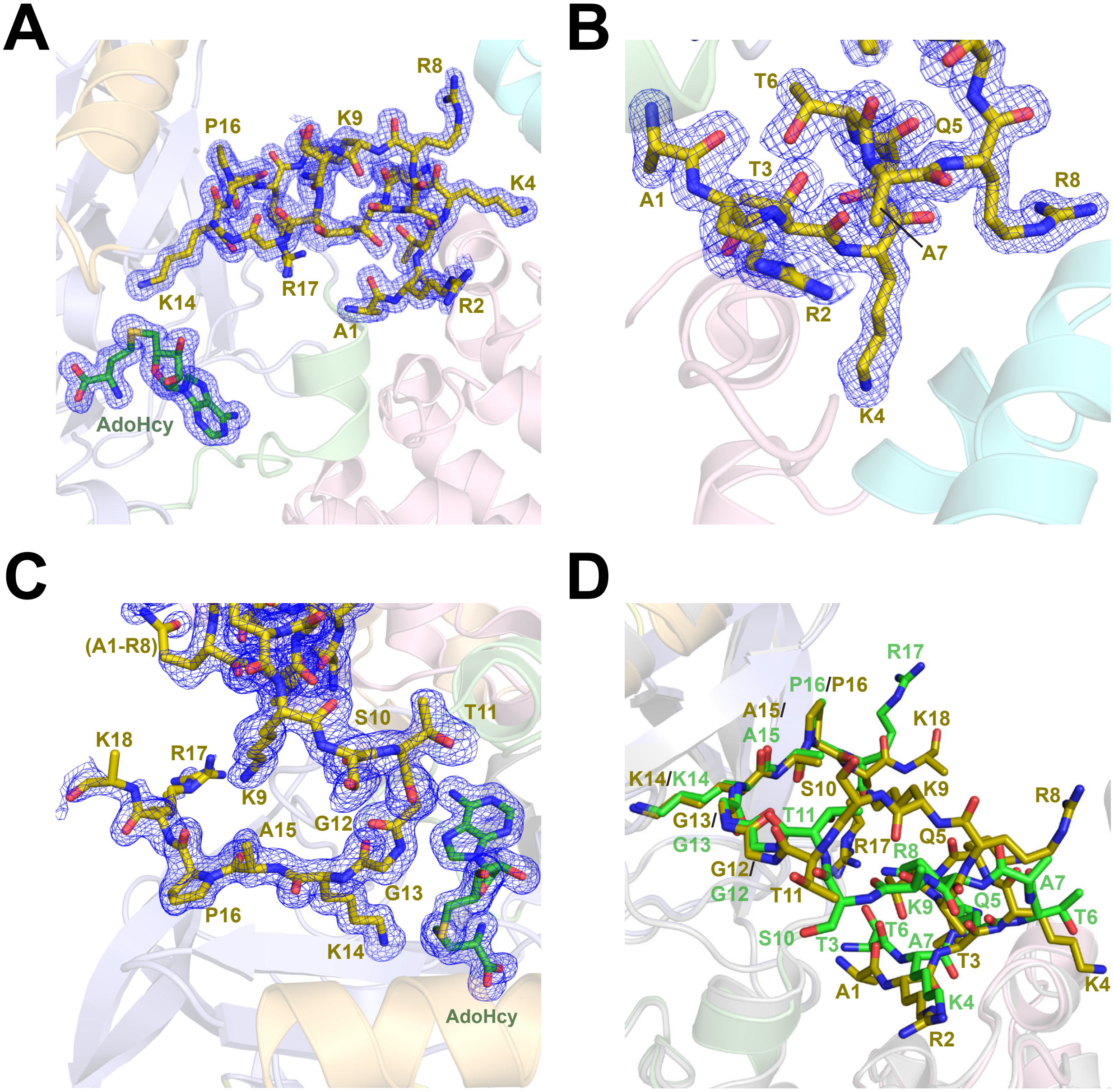

**Figure S4: Binding of histone H3 peptides and AdoHcy to LegAS4. A.** Simulated-annealing omit maps for the histone H3.1$_{[1-21]}$ peptide (gold carbon atoms) and AdoHcy (green carbon atoms) bound to LegAS4 (colored as Figure 3A) from the structure of the LegAS4/AdoHcy/H3$_{[1-21]}$ complex. The 2mFo-DFc electron density of the omit map is contoured at 1.0 sigma. **B, C.** Zoomed in view of the omit map for the histone H3 residues bound to the Ankyrin repeat domain (**B**) and the SET domain (**C**). **D.** Structural alignment of the LegAS4/AdoHcy/H3$_{[1-21]}$ complex (colored as in panel **A**) and the previously determined LegAS4/AdoHcy/H3$_{[3-17]}$ complex (PDB 8SR6, light gray cartoon) with the histone H3 carbon atoms depicted in gold and light green, respectively.

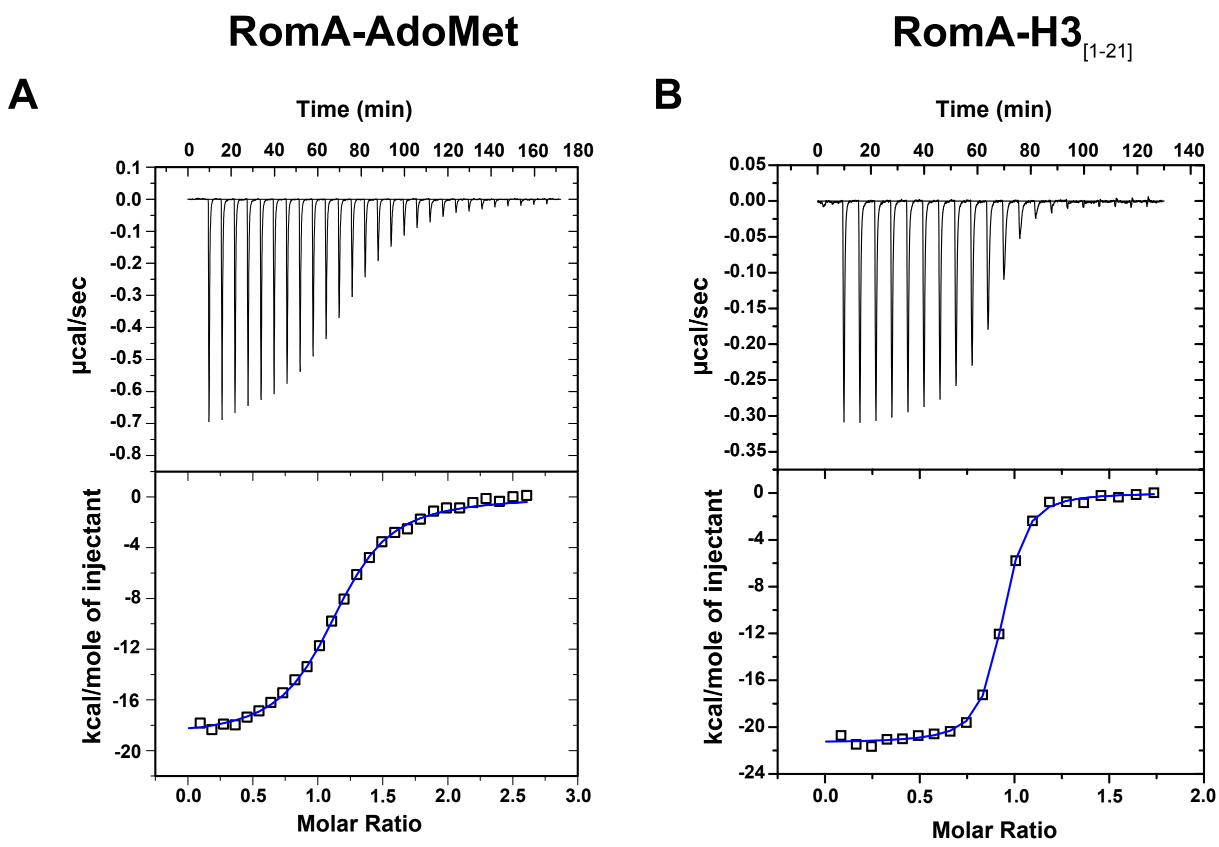

**C**

| Protein-Ligand | $K_D$ (nM) | STD |
|---|---|---|
| RomA-AdoMet | 424.4 | ± 20.3 |
| RomA-$H3_{[1-21]}$ | 19.7 | ± 3.5 |

**Figure S5: ITC analysis of substrate binding to RomA. A., B.** Representative titration binding experiments (upper panels) and binding isotherms with the fitted binding curves (lower panels) for the association of RomA with AdoMet (**A**) and the $H3_{[1-21]}$ peptide (**B**). **C.** Summary of equilibrium dissociation constants ($K_D$) for the binding of RomA to its substrates. Mean $K_D$ values and their standard deviations (STD) are reported from triplicate measurements.

**A**

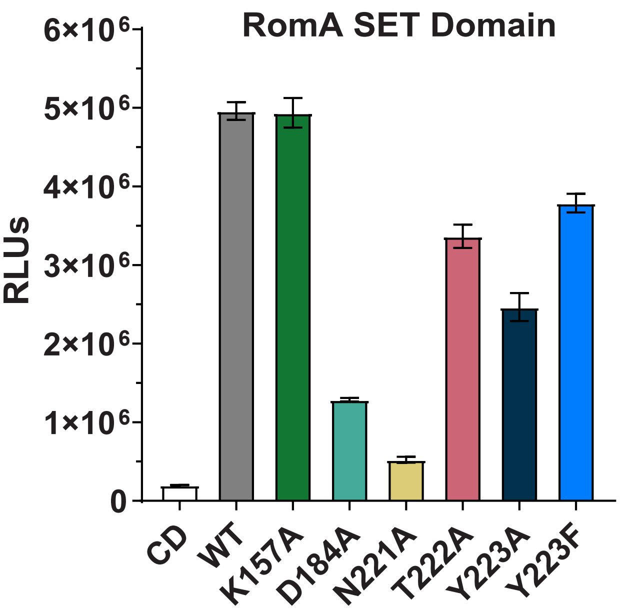

**B**

| | -3 | -2 | -1 | 0 | +1 | +2 | +3 |
|---|---|---|---|---|---|---|---|
| **H3K4** | A | R | T | K | Q | T | A |
| **H3K9** | T | A | R | K | S | T | G |
| **H3K14** | T | G | G | K | A | P | R |
| **H3K27** | A | A | R | K | S | T | P |
| **H3K36** | G | G | V | K | K | P | H |
| **H4K20** | R | H | K | K | V | L | R |

**Figure S6: H3K14 recognition by the SET domain. A.** Second replicates of the luminescent methyltransferase assays shown in Figure 4B. **B.** Sequence alignment of H3K14 ± three residues with the major lysine methylation sites in histone H3 and H4 illustrates that only the H3K14 site possesses the LpHKMT sequence recognition motif G-K-X-(P>A) motif (where K is the lysine substrate and X is any amino acid).

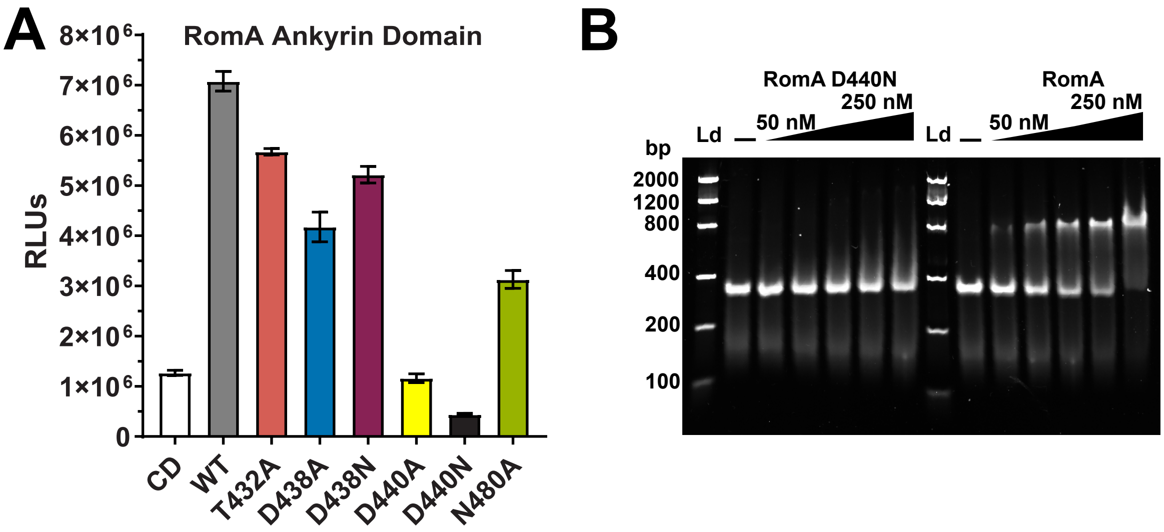

**Figure S7: Recognition of H3 N-terminal tail by the Ankyrin repeat domain. A.** Second replicates of luminescent methyltransferase assays in Figure 5D. **B.** Second replicate of the EMSAs in Figure 5F.

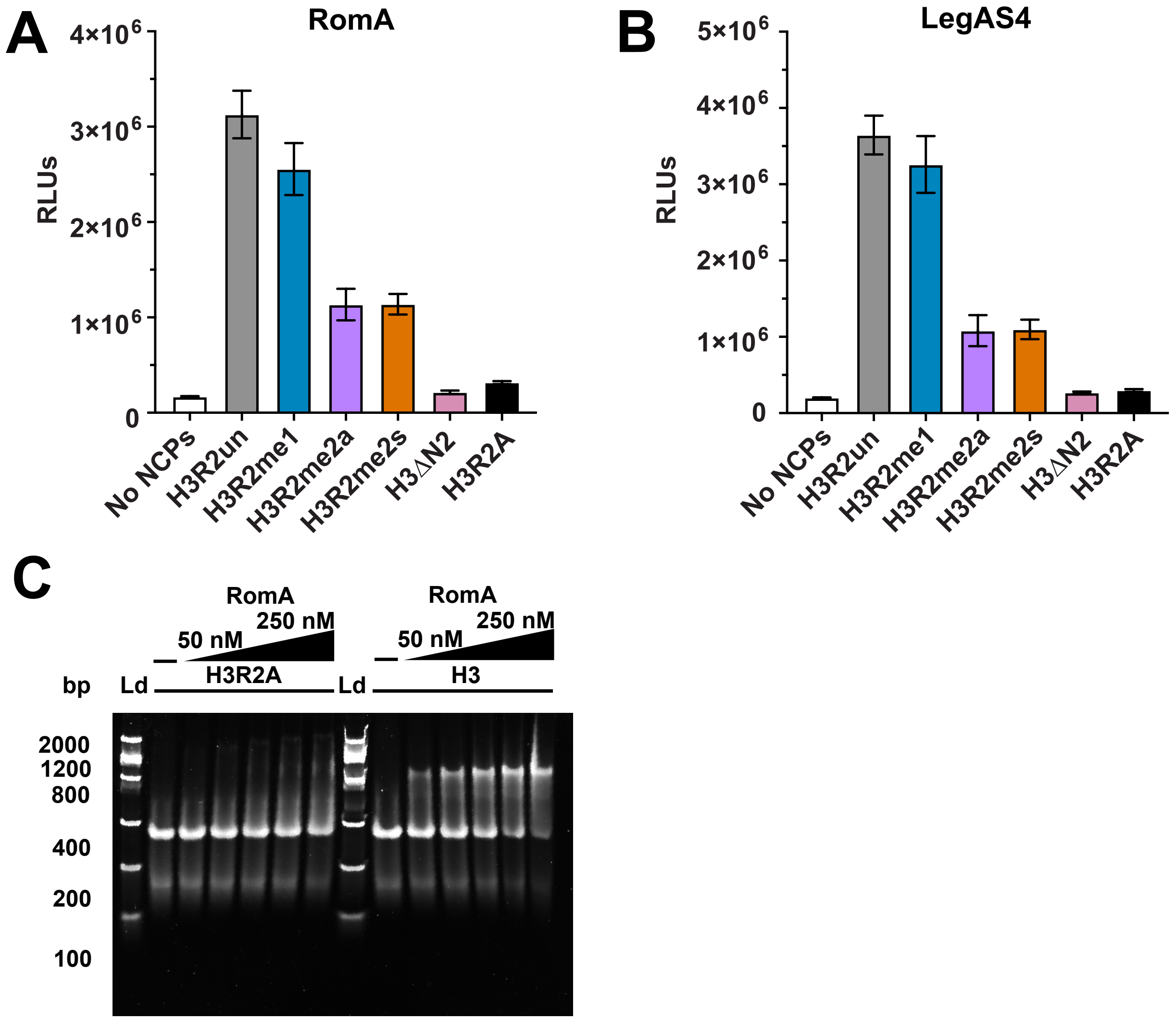

**Figure S8: Recognition of H3R2 methylation by the LpHKMTs. A, B.** Second replicate of the luminescent methyltransferase assays in Figure 6B, 6C. **C.** Second replicate of the EMSAs in Figure 6D.

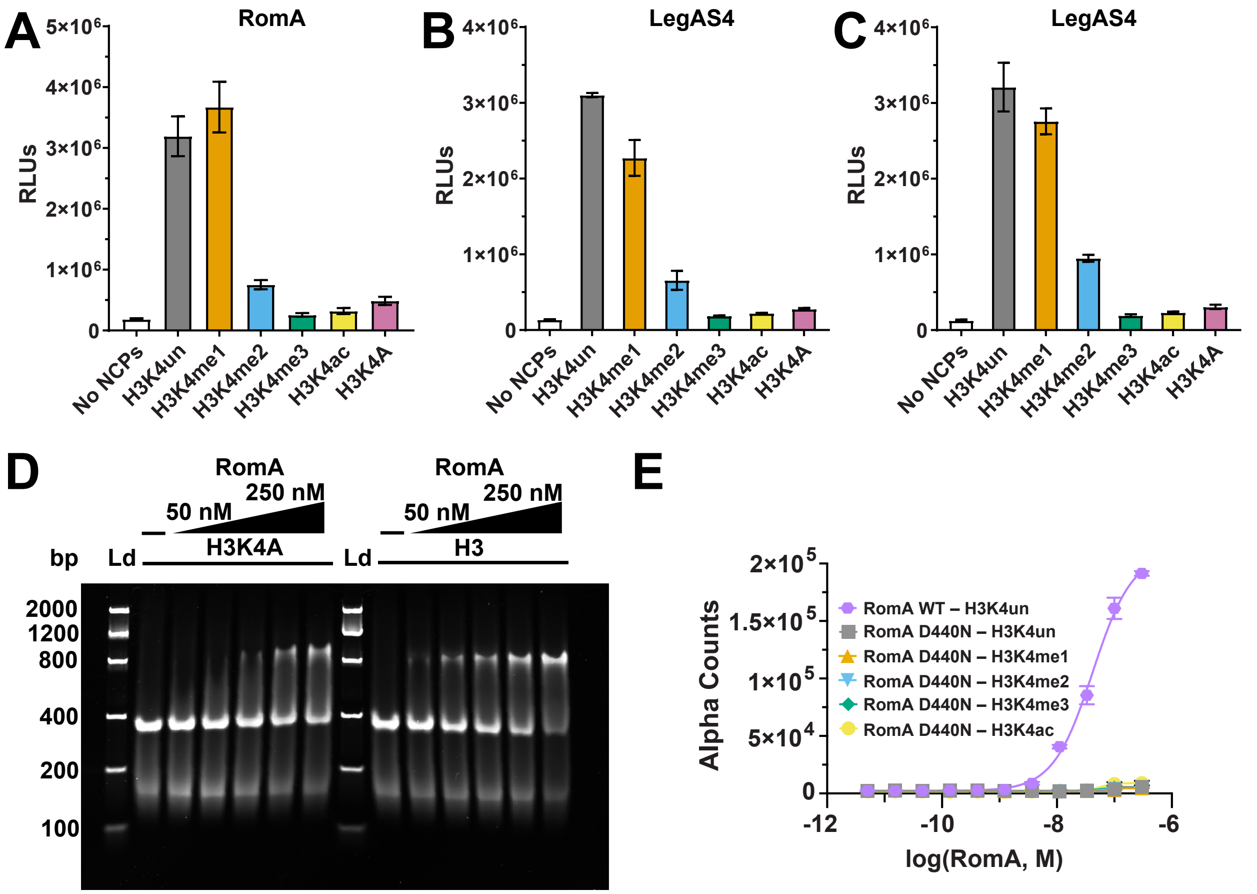

**Figure S9: Reading of H3K4 modifications by the Ankyrin repeat domain. A.** Second replicate of the luminescent methyltransferase assays in Figure 8B. **B, C.** Independent replicates of methyltransferase assays of LegAS4 using NCPs with ([H3K4]$_2$) PTMs or ([H3.1K4A]$_2$) as substrates (Table S1). **D.** Second replicate of the EMSAs in Figure 8C. **E.** Captify-Alpha assays with RomA D440N and NCPs bearing a diverse PTM landscape at H3K4. The engagement of RomA WT with ([H3K4un]$_2$) is a positive control.

### SUPPLEMENTARY METHODS:

#### Bacterial strains and culture conditions

The parent strain for our studies was *Legionella pneumophila* Lp02, a thymidine auxotroph laboratory strain derived from the clinical isolate *L. pneumophila* Philadelphia-1. All strains analyzed were also flagellin-deficient (Lp02 Δ*flaA::kan*) to mitigate cytotoxic effects on macrophages as reported previously [1,3]. *L. pneumophila* strains were cultured in ACES-buffered yeast extract (AYE) broth at pH 6.9 supplemented with 0.1 mg/ml thymidine, 0.4 mg/ml L-cysteine, and 0.135 mg/ml ferric nitrate (AYET medium) or on solid medium containing 15 g/liter agar and 2 g/liter activated charcoal (CYET agar) [2]. AYET medium and CYET agar were supplemented where necessary with 5 µg/ml chloramphenicol or 25 µg/ml kanamycin.

Prior to an experiment, bacterial strains from glycerol stocks maintained at -80 °C were struck onto CYET agar and incubated at 37 °C for 3 – 4 days. Fresh colonies (< 1-week old) were cultured in AYET medium overnight at 37 °C, and then sub-cultured in AYET medium overnight to exponential phase ($OD_{600}$ of 0.5 – 2.0) (for strain construction) or post-exponential phase ($OD_{600}$ of 3.5 – 4.5) (for macrophage infections).

#### *legAS4* deletion mutant construction

The Δ*legAS4* mutant strain was constructed as previously [3] from the parent strain *L. pneumophila* Lp02 Δ*flaA::kan* (Table S6). In brief, a chloramphenicol resistance cassette (chloramphenicol acetyltransferase; *cat*) flanked by Flp recognition sites was introduced at the *legAS4* locus. After Flp-mediated excision of *cat*, a clean deletion of the *legAS4* locus was verified by sequencing and loss of antibiotic resistance.

The *legAS4* ~1,600-bp genomic locus (*lpg1718*) with ~500-bp of flanking DNA was amplified from the Lp02 Δ*flaA::kan* genome using primer pair 192/193, then ligated using pGEM-T Easy kit to generate pGEM-*legAS4* (Table S7, S8). The ~1 kb *cat* cassette from pKD3 [4] was amplified with ~35-bp of DNA homologous to the regions 3' and 5' of *legAS4* using primer pair 185/186. A recombinant allele, *legAS4::cat*, was generated by λ-red recombineering in *Escherichia coli* DY330 [5] and amplified by PCR using primer pair 192/193 (Table S7).

The recombinant allele was introduced and integrated into Lp02 Δ*flaA::kan* by natural transformation and homologous recombination. The *cat* cassette was excised by Flp recombinase following transformation of the isolated deletion mutant with pBSFlp as previously [6]. The unmarked deletion of *legAS4* was verified by DNA sequencing and loss of antibiotic resistance.

**Complementation tests**

To evaluate genetic complementation, the wild type *legAS4* gene and its native promoter were inserted into a neutral site in the Lp02 Δ*legAS4* chromosome between *lpg2528* and *lpg2529* [7]. We first constructed plasmid pSU-pZLCm from pSU-pZLKm [7] (Table S9). To do so, the *cat* cassette from pKD3 [4] was amplified with Gibson primer pair 268/270 (Table S7). The PCR product was cloned into a linearized pSU-pZLKm backbone – amplified with Gibson primer pair 269/271 – by Gibson Assembly (NEB), producing pSU-pZLCm. Next, *legAS4* with ~500-bp of flanking DNA was amplified from Lp02 genomic DNA with primer pair 127bis/128 and ligated into the BamHI and KpnI sites of pSU-pZLCm (Table S6-7). Wild-type *romA* was similarly cloned into pSU-pZLCm after amplification from *L. pneumophila* Paris genomic DNA with primer pair 129bis/128 (Table S7).

To analyze a catalytically defective LegAS4 protein, a *legAS4* gene with six point mutations targeting the S-adenosylmethionine (AdoMet) binding pocket was generated by successive site-directed mutagenesis PCR on pSU-pZLCm-*legAS4* with respective primer pairs 202/203, 204/205, 206/207, 246/247, 284/285, and 286/287 (Table S7) [8]. Single HA epitope tags (YPYDVPDYA) were inserted at the C-terminus of LegAS4 and RomA by one-step site-directed insertion mutagenesis PCR on pSU-pZLCm-*legAS4* or -*romA* constructs with respective primer pairs 260/261 and 262/263 (Table S7) [8].

After transformation of the Lp02 Δ*legAS4* Δ*flaA::kan* mutant strain, each plasmid integrated into the intergenic region between *lpg2528* and *lpg2529* to generate deletion mutant strains carrying a chromosomal allele of either *legAS4*, *romA*, or catalytically deficient *legAS4*. To generate a negative control strain, the Lp02 Δ*legAS4* Δ*flaA::kan* strain was transformed with the pSU-pZLCm vector. Each strain construction was verified by growth on CYET + kan/cam and PCR prior to testing genetic complementation by quantifying intracellular growth of *L. pneumophila*.

**Isolation and culturing of bone marrow-derived mouse macrophages**

Macrophages were derived from the bone marrow of female A/J mice obtained from The Jackson Laboratory (between 6-13 weeks of age) as previously [9]. Following euthanasia by $CO_2$ overdose and cervical dislocation, both femurs and tibias were dissected and cleared of adherent tissue. The ends of the bones were cut off and the bone marrow flushed into a sterile petri dish with 5 mL RPMI 1640 (Table S8). Cells were suspended by vigorous pipetting and vortexing and washed once by centrifugation at 200 x g for 10 min. Cells were cultured in non-treated petri dishes in RPMI 1640 containing 20% heat-inactivated fetal bovine serum (Table S8), 2 mM

GlutaMax (Table S8), 50 ng/mL recombinant human M-CSF (Table S8), 100 U/ml penicillin-streptomycin (Table S8), and 35 mM β-mercaptoethanol (Table S8). After 6 to 7 days of incubation at 37 °C in a humidified chamber containing 5% $CO_2$, differentiated macrophages were washed with ice-cold phosphate-buffered saline (Table S8), chilled on ice, and collected from petri dishes.

Each mouse yielded between 1 – 2 x $10^8$ cells, which were divided into aliquots and frozen in full RPMI media with recombinant human M-CSF, as described above. Cells were slow frozen in an isopropanol freezing container at -80 °C and then were stored in liquid nitrogen. Cell aliquots were thawed and plated in tissue culture wells in full RPMI medium with recombinant human M-CSF for 3 – 4 hours, then cultured for an additional 18 – 24 hours in RPMI with 10% FBS prior to starting an experiment.

**Intracellular bacterial growth**

Mouse macrophages cultured as described above were plated at 2.5 x $10^5$ cells per well of a 24-well tissue culture plate and incubated overnight (18 – 24 hours) prior to infection as previously [10]. Post-exponential phase cultures of *L. pneumophila* strains were diluted in infection media (RPMI 1640 with 10% FBS and 100μg/ml thymidine to support *L. pneumophila* growth) and added to the macrophages at a Multiplicity of Infection (MOI) of 1 for 2 hours at 37 °C in a humidified chamber containing 5% $CO_2$. Extracellular bacteria were then removed by washing the macrophage monolayers three times with RPMI at 37 °C, and fresh infection media was added to each well. At 2, 24, 48, and 72 hours, culture supernatants were collected, cells lysed with 0.5% saponin in PBS, and appropriate dilutions of triplicate samples plated on CYET agar. The total number of colony-forming units (CFU) per well was determined by enumerating CFU after 3 – 4 days of incubation at 37 °C. Results are expressed as the fold increase of CFU recovered from the cell lysates at 24-hour interval time points compared to the 2-hour time point for each *L. pneumophila* strain analyzed.

**Immunoblot analysis**

Mouse macrophages cultured as described above were plated at a density of $10^6$ cells per well of a 6-well tissue culture plate and infected after overnight incubation (18 – 24 hours) as previously [10]. Post-exponential phase cultures of *L. pneumophila* strains were diluted in infection media (RPMI 1640 with 10% FBS and 100μg/ml thymidine to support *L. pneumophila* growth) and added to the macrophages at an MOI of 5 for 1 hour at 37 °C in a humidified chamber containing 5% $CO_2$. Extracellular bacteria were removed by washing the macrophage monolayers three times with RPMI at 37 °C, and fresh infection media was added to each well. At 1, 2, 4, and 6 hours, cells were lysed and collected in lysis buffer (50 mM Tris pH 7.5, 50 mM NaCl, 10% glycerol, 0.5% Triton X-100, 0.1% NP-40, 1X HALT Protease inhibitor without EDTA [Table S8]). Total protein in each lysate sample was quantified by BCA Protein assay (Table S8).

Cell lysates were boiled in Laemmli Buffer (Table S8) for 10 min at 95 °C and needle sheared. 5 µg of total protein per sample was resolved by SDS-PAGE (Table S8) and transferred to polyvinylidene difluoride (PVDF) membranes (Table S8) by electrophoresis at 4 °C in 20% methanol, 192 mM glycine, 25 mM Tris at 80 V for 90 min. The membranes were blocked in Tris-buffered saline with 0.05% Tween-20 (TBST) containing 5% nonfat milk (blocking buffer) and incubated at 4 °C overnight with anti-H3 rabbit monoclonal antibody diluted 1:1000 in blocking buffer or anti-H3K14me2 mouse monoclonal antibody diluted 1:2000 in blocking buffer (Table S10). Membranes were washed 5 times in TBST and incubated for 1 hour at room temperature with secondary goat anti-mouse or anti-rabbit antibody conjugated to horseradish peroxidase diluted 1:2000 in TBST. Membranes were washed 5 times in TBST and developed with Pierce ECL Western Blotting Substrate (Table S8). Immunoblots were imaged with the Bio-Rad ChemiDoc system.

Membranes were incubated overnight at 4 °C with anti-β-actin HRP-conjugated antibody (Table S10) diluted 1:4000 in blocking buffer and developed and imaged as described above as a protein loading control. Recombinant histone H3.1 (0.1 µg) (Table S8) and histone H3 (0.1 µg) methylated *in vitro* by LegAS4 (25 nM LegAS4, 5 µM histone H3.1, 100 µM S-(5'-Adenosyl)-L-methionine chloride (AdoMet, [Table S8], 50 mM Tris pH 8.0, 100 mM KCl, 1 mM DTT; 30 min reaction at room temperature, quenched with Laemmli buffer) were included in the immunoblot analysis as controls for H3K14 methylation.

### Nucleosome reconstitution

Histone mutagenesis and expression

Histone H2A, H2B, H3.1, and H4 plasmids were kindly provided by Dr. Uhn-Soo Cho, from the University of Michigan. Histone H2A was cloned into vector pMCSG7 which contains a hexahistidine (6xHis) tag and TEV protease site [11]. Histone H2B, H3.1, and H4 were cloned into the pET3a vectors [11]. Point mutations for H3.1R2A, H3.1K4A, and H3.1K14M were introduced via site directed mutagenesis (Table S7). Cys110Ala (C110A) on *X. laevis* histone H3.1 wild type and mutants were introduced using the same method to eliminate disulfide bond formation [12, 13, 14].

Wild type *X. laevis* core histones H2A, H2B, and H4 along with H3.1 C110A, H3.1R2A C110A, H3.1K4A C110A, and H3.1K14M C110A were individually expressed and purified as previously [11]. Plasmids were freshly transformed into *E. coli* Rosetta2-DE3 cells with the addition of the pRARE2 gene on LB plates with antibiotics (100 μg/mL ampicillin and 25 μg/mL chloramphenicol) (Table S8). Several transformed colonies were used to inoculate 125 mL of terrific broth starter culture (Table S8). The starter culture was grown at 37 °C until $OD_{600}$ reached a value >0.5. Starter culture was inoculated into 500 mL of LB broth with antibiotics (100 μg/mL ampicillin and 25 μg/mL chloramphenicol) supplemented with glucose to a final concentration of 0.2% v/v (Table S8). The culture was incubated at 37 °C until $OD_{600}$ reached a range of 0.6 – 0.8. Cells were induced at a final concentration of 0.5 mM IPTG at 37 °C for 2 – 3 hours. Culture temperature was turned down to 18 °C and remained shaking overnight. Cells were harvested by centrifugation (5,775 x g) for 30 minutes, resuspended in 50 mM Tris-HCl pH 7.4, 100 mM NaCl, 1 mM EDTA, 1 mM DTT, then flash frozen in liquid nitrogen to be stored at -80 °C.

| **Histone Octamer Refolding and Purification Buffers** | |
|---|---|
| **Buffer Name** | **Buffer Components** |
| Histone Buffer Wash TW (TW Buffer) | 50 mM Tris-HCl pH 7.4, 1 mM EDTA, 100 mM NaCl, 1 mM DTT, 1% v/v Triton X-100, and 2.5 mL resuspended Sigma-Aldrich bacterial protease inhibitor cocktail |
| Histone Buffer Wash T (T Buffer) | 50 mM Tris-HCl pH 7.4, 1 mM EDTA, 100 mM NaCl, 1 mM DTT, and 2.5 mL resuspended Sigma-Aldrich bacterial protease inhibitor cocktail |
| Inclusion Body Buffer (IB Buffer) | 8M Guanidine-HCl, 20 mM Sodium Acetate pH 5.2, 10 mM DTT |
| Histone Octamer Dialysis Buffer (Dialysis Buffer) | 20 mM Tris pH 8.0, 2M NaCl, 2 mM βME |
| Histone Octamer Equilibration Buffer (EQ Buffer) | 2M NaCl, 20 mM Tris pH 8.0, 10 mM Imidazole, 2 mM βME |
| Histone Octamer Elution Buffer (E Buffer) | 2M NaCl, 20 mM Tris pH 8.0, 500 mM Imidazole, 2 mM βME |
| Histone Octamer Gel Filtration Buffer (GF Buffer) | 2M NaCl, 20 mM Tris pH 7.5, 1 mM DTT |
| Histone Octamer Storage Buffer (S Buffer) | 20 mM Tris pH 7.5, 2M NaCl, 1 mM EDTA, 1 mM DTT, 50% v/v glycerol |

Individual histones pellets were thawed completely and combined in an ~ equal ratio based on expression with an excess of H2A and H2B to avoid the formation of hexamers (lacking one H2A–H2B dimer)[15]. Combined cell pellets were centrifuged (37,157 x g) for 20 minutes at 4 °C and the supernatant was discarded. 25 mL of TW Buffer was used to resuspend and homogenize the pellets (Table S8). Cells were sonicated in the presence of lysozyme. Sonicated cells were centrifuged (37,157 x g) for 20 minutes at 4 °C and supernatant discarded. 25 mL of TW Buffer was used to resuspend and homogenize the pellet. Cells were then centrifuged (37,157 x g) for 20 minutes at 4 °C and supernatant discarded. Pellets were resuspended and homogenized in 20 mL of T Buffer. Cells were centrifuged (37,157 x g) for 20 minutes at 4 °C.

Pellets were again resuspended and homogenized in 20 mL of T Buffer and centrifuged (37,157 x g). Cell pellets were flash frozen in liquid nitrogen and stored at -80 °C.

Histone octamer refolding and purification

Histone octamers were prepared using the one-pot method[11]. Inclusion body cell pellets were thawed in DMSO. 10 mL of IB Buffer was added to each pellet and minced. Cell pellets were stirred at room temperature for 1 hour. Inclusion bodies were then centrifuged (38,724 x g) for 40 minutes at 4 °C. Supernatant was set aside on ice. 5 mL of IB Buffer was added, pellets were minced and stirred at room temperature for 30 minutes, and inclusion bodies again collected by centrifugation. Supernatants were pooled and dialyzed overnight at 4 °C in 4L of Dialysis Buffer (Table S8).

Dialysis bags were transferred to 4L fresh Dialysis Buffer three times at 8-hour intervals. Dialysis contents were centrifuged (185,511 x g) for 40 minutes at 4 °C, 0.45 μm filtered, and applied onto 4 mL Ni-NTA column (Table S8). The column was washed with EQ Buffer, and histone octamers were eluted using a 10 mM – 500 mM Imidazole linear gradient in E buffer. Peak fractions identified by SDS-PAGE were pooled and TEV Protease S219V added to cleave the 6xHis Affinity tag. TEV Protease S219V was expressed and purified as previously[16]. Fractions were dialyzed overnight at 4 °C against 4L of EQ Buffer. Cleaved protein was incubated with 6 mL Ni-NTA resin for 30 minutes at 4 °C, and the flow through collected after a gravity column. The column was washed with EQ Buffer to remove residual cleaved protein, the pool concentrated, applied to Superdex 200, and eluted by step-gradient using GF Buffer.

Peak fractions were analyzed on SDS-PAGE, then pooled and concentrated. Histone octamers (in 10 kDa MWCO dialysis cassette) were dialyzed overnight at 4 °C in S Buffer, aliquoted, flash frozen in liquid nitrogen, and stored at -80 °C [15].

Preparation and purification of 601 nucleosomal DNA

Preparation of 147-bp 601 nucleosomal DNA oligonucleotide (601 DNA) was as previously[11, 17, 18], with modifications. The 601 DNA plasmid was transformed into *E. coli* DH5α (Table S6). After transformation, flasks were inoculated and grown with antibiotics (100 µg/mL ampicillin) overnight at 37 °C.

Cell pellets from 2.5 L of cell culture were collected and plasmid DNA isolated using a Plasmid Giga-prep kit (Table S8) and quantified by absorbance at 260 nm. The insert-containing construct was digested with excess EcoRV-HF restriction endonuclease (Table S8) for 16 hours at 37 °C to separate the 601 DNA fragment from the vector backbone. The endonuclease was heat-inactivated at 65 °C for 20 minutes, and DNA analyzed on a 1% agarose gel to verify complete digestion. 601 DNA was separated from the plasmid backbone using a Source 15Q ion-exchange column (Table S8) and eluted by a linear salt gradient (0 – 1M KCl) in 10 mM Tris pH 7.5, 1 mM EDTA. Peak fractions were analyzed by agarose gel electrophoresis, pooled, and dialyzed overnight into Milli-Q water at 4 °C using 2kDa MWCO dialysis cassette (Table S8). 601 DNA concentration was determined using a NanoDrop spectrophotometer at 260 nm before lyophilization and storage at -20 °C.

Nucleosome assembly

*X. laevis* histone octamers (H3.1, H3.1R2A, H3.1K4A, and H3.1K14M [all C110A]) in 50% glycerol v/v (Table S1) were thawed and dialyzed (2M KCl, 20 mM Tris pH 7.5, 1 mM EDTA, 1 mM DTT) to remove glycerol using a 10kDa MWCO dialysis cassette (Table S8). Histone octamer concentration was then measured by absorbance at 280 nm using the NanoDrop spectrophotometer.

Nucleosome core particle (NCP) complexes were assembled using 601 DNA as previously [11, 17, 19, 20]. 601 DNA was resuspended in water, quantified at 260 nm, and then mixed with an equal volume of 4M KCl. Histone octamers were added directly to the 2M KCl-601 DNA solution (1:1.7 mass ratio, respectively) and then placed in a 3.5 kDa MWCO cassette (Table S8). Assembly of the NCPs was achieved by dialyzing the histone octamer-601 DNA mixture from high-salt buffer (2M KCl, 20 mM Tris-HCl pH 7.5, 1 mM EDTA, 1 mM DTT) into no-salt buffer (20 mM Tris-HCl pH 7.5, 1 mM EDTA, 1 mM DTT) using a peristaltic pump for 3 days at 4 °C.

Reconstituted NCPs were centrifuged (15,600 x g) at 4 °C to separate precipitated material. The supernatant was heat shifted for 30 minutes at 55 °C (for DNA repositioning), then placed on ice for 10 minutes. NCPs were concentrated using a 50 kDa MWCO and validated by 5% TBE gel electrophoresis using ethidium bromide, followed by Coomassie Blue staining (Table S8). NCPs were concentrated >10 µM as determined using a NanoDrop spectrophotometer at 260 nm, and glycerol was added to a final concentration of 20% v/v. NCPs were aliquoted, flash-frozen in liquid nitrogen, and stored at -80 °C [15].

**Cloning & mutagenesis of the LpHKMTs**

Synthetic *legAS4* and *romA* genes codon-optimized for expression in *E. coli* were synthesized by GeneArt Gene Synthesis Service (Thermo Fisher Scientific). Each LpHKMTs gene was cloned into BamHI and XhoI sites of ppSUMO (Table S9) for expression as N-terminal 6xHis-tagged SUMO fusion proteins [21]. Deletion and site-directed mutagenesis were performed by PCR using partially overlapping primers with Q5 High-Fidelity Polymerase (Table S7, S8) [8]. For the Captify™-Alpha assays, a Strep-SUMO RomA construct possessing a C-terminal 6xHis-tag was generated by substituting the DNA sequence encoding the N-terminal 6xHis-tag in the ppSUMO-RomA plasmid with the Strep-tag encoding sequence by deletion and insertion mutagenesis [8]. The C-terminal 6xHis-tag was incorporated at the 3'-end of the *romA* gene by deleting the native stop codon at the 3'-end of the protein coding sequence, permitting translational read through to the 3'-6xHis-tag and stop codon encoded after the multicloning site in the plasmid.

**Expression and purification of the LpHKMTs**

Recombinant N-terminal 6xHis-tagged SUMO fusion proteins of LegAS4, RomA, and RomA mutants were expressed in *E. coli* Rosetta2-DE3 cells in 2xYT medium and induced with 0.5 mM IPTG overnight at 18 °C (Table S6, S7, S8). After cell lysis and centrifugation, fusion proteins were purified by TALON Superflow (Table S8) and eluted with a linear gradient of imidazole (0 – 300 mM) in 50 mM Sodium Phosphate pH 7.5, 500 mM NaCl, and 5.0 mM β-mercaptoethanol (βME) using a Bio-Rad NGC chromatography system. Elution fractions were analyzed by SDS-PAGE, and the peak pooled and dialyzed overnight against 25 mM Sodium Phosphate pH 7.5, 150 mM NaCl, and 5.0 mM βME in the presence of 6xHis-tagged SUMO protease Ulp1. Dialyzed protein was purified using TALON Superflow batch step to remove the 6xHis-tagged SUMO and Ulp1, as well as any fusion protein not cleaved by Ulp1. Protein recovered from the TALON Superflow batch step was concentrated and purified by gel filtration chromatography using a Superdex 200 column (Table S8). WT enzyme and mutants displayed the same elution volume. Peak fractions eluted from the column were analyzed by SDS-PAGE and the purest fractions pooled and concentrated. The concentration of the purified protein was determined by its absorbance using its 280 nm extinction coefficient calculated by the ProtCalc tool at the JustBio website (www.justbio.com). Concentrated protein was aliquoted into microfuge tubes, flash frozen in liquid nitrogen, and stored at -80 °C.

Captify™-Alpha nucleosome binding assays required 6xHis-tagged RomA constructs [22]. Strep-SUMO RomA proteins with an uncleavable C-terminal 6xHis-tag were purified by affinity chromatography using a Strep-Tactin Superflow Plus column (Table S8) and eluted by a step gradient from 0 – 2.5 mM desthiobiotin in 50 mM Sodium Phosphate pH 7.5, 500 mM NaCl, and 5.0 mM βME. Fractions from the Strep-Tactin column were analyzed by SDS-PAGE, and peak fractions pooled and dialyzed overnight in the presence of Ulp1 to digest the SUMO-RomA fusion protein, as described above. The following day, the resulting C-terminal 6xHis-tagged RomA protein was purified by TALON Superflow and Superdex 200 gel filtration chromatography, as described above. Peak fractions from the Superdex 200 column were analyzed by SDS-PAGE, and the purest fractions pooled and concentrated. Protein concentration was determined by absorbance at 280 nm, aliquoted, frozen in liquid nitrogen, and stored at -80 °C.

For isothermal titration (ITC) experiments to quantify the binding of the RomA Δ1-70 construct to S-adenosylmethionine (AdoMet) and to a histone $H3_{[1-21]}$ peptide, the enzyme was denatured and refolded to remove AdoMet bound during expression in *E. coli*. The 6xHis-SUMO-RomA Δ1-70 fusion protein was expressed, as described above. After cell lysis and centrifugation, fusion proteins were purified by TALON Superflow (Table S8) and eluted with a linear gradient of imidazole (0 – 300 mM) in 50 mM Sodium Phosphate pH 7.5, 500 mM NaCl, 6.0 M Guanidinium Chloride, and 5.0 mM β-mercaptoethanol (βME) to denature the enzyme and remove the bound AdoMet using a Bio-Rad NGC chromatography system. Elution fractions were analyzed by SDS-PAGE, the peak pooled, and dialyzed overnight against 50 mM Sodium Phosphate pH 7.5, 500 mM NaCl, and 5.0 mM βME in the presence of 6xHis-tagged SUMO protease Ulp1. Following cleavage by Ulp1, RomA Δ1-70 was further purified by Talon Superflow batch step and Superdex 200 gel filtration chromatography, as described above.

**Isothermal titration calorimetry**

The binding affinity of RomA for its substrates was measured using ITC. Titration experiments were performed in ITC solution (20 mM Sodium Phosphate pH 7.5 and 125 mM KCl) at 20 °C using a MicroCal VP-ITC calorimeter (Malvern Panalytical). AdoMet (125 μM) was titrated into a solution of the RomA Δ1-70 construct (10 μM), whereas histone $H3_{[1-21]}$ peptide (40 μM) was titrated into a solution of RomA Δ1-70 (5.0 μM). A single-site binding model was fit to the binding isotherm to determine the equilibrium dissociation constant ($K_D$) using Origin 7.0 (OriginLab Corp.). Titrations were performed in triplicate with binding stoichiometries (N values) between 0.8 – 1.0.

**Electrophoretic mobility shifts assays**

Binding reactions were assembled on ice using a stepwise concentration gradient of RomA and LegAS4 (0 – 250 nM with 50 nM increments) in EMSA binding buffer (20 mM Tris pH 7.5, 100 mM KCl, 1 mM TCEP, 2 mM EDTA, 0.1 mg/mL recombinant Albumin 0.05% NP-40, 10% Glycerol) (Table S8). 5% TBE PAGE gels were pre-run for 1 hour at 100V at room temperature in 0.2X TBE buffer (Table S8). Binding reactions were initiated by adding NCPs to a final concentration of 25 nM and incubated for 45 minutes on ice. Prior to loading, 0.2X TBE buffer was replaced with a fresh cold 0.2X TBE buffer and placed at 4 °C. Binding reactions were then loaded on the 5% TBE gels and electrophoresed for 60V at 4 °C on ice for 3.5 hours. After completion, gels were stained using 0.5X SYBR Gold solution (prepared as a 1:20,000 dilution with filtered 0.2X TBE buffer [Table S8]) for 10 minutes at room temperature with gentle rocking. After staining, gels were washed once with 0.2X TBE buffer and visualized using a Bio-Rad ChemiDoc System.

**Luminescent methyltransferase-glo Assays**

Methyltransferase-Glo™ assays reactions (Table S8) were performed in white 384-well plates (Table S8) using 2X Enzyme-NCP mixtures containing the MTase-Glo™ Reagent (Table S8), 10 nM LpHKMT, 2 µM nucleosome, 2 mM DTT, and 100 mM KCl, in a total volume of 5 µl [23]. Assays were initiated by adding 5 µl of 200 µM AdoMet (Table S8) at room temperature. After 30 minutes, 10 µl MTase-Glo™ Detection Solution (Table S8) was added, mixed thoroughly, and incubated for 30 minutes at room temperature. Luminescence was measured using a Promega GloMax plate reader. Bar graphs were generated using GraphPad Prism [24].

**Fully defined nucleosomes**

This study uses a histone and nucleosome nomenclature recently devised for accurate scientific communication in the chromatin field [25]. Histone peptides are indicated by subscripted brackets identifying the start and end, followed by any PTMs in order (*e.g.*, $H3_{[1-12]}$K4me3). Any distinguishing histones in a fully defined semi-synthetic nucleosome are indicated, with other positions not denoted understood to be unmodified major histones (as in ($[H3K4me3]_2$)). Designer nucleosomes (dNucs) were made using native chemical ligation to yield full-length 'scarless' histones; versaNucs were made by enzymatic ligation of histone $H3_{[1-31]}$A29L peptides with a designation of interest to a H3 tailless nucleosome precursor (*e.g.,* ($[H3\Delta N32]_2$); *EpiCypher* 16-0016) [26, 27].

**Captify™-Alpha binding assays**

The assay previously known as dCypher™ is now named Captify™, with no distinction in how the assay is performed or its capabilities. Assays on the no-wash Alpha (amplified luminescence proximity homogeneous assay) platform (*PerkinElmer*, *Revvity*) to examine the interaction of RomA-6xHis (the Query) with biotinylated PTM-defined nucleosomes (the targets) were as previously [22, 27, 28], with all bead handling and incubation performed under subdued lighting. Nucleosome assay buffer was (20 mM Tris, pH 7.5, 100–250 mM NaCl, 0.01% BSA, 0.01% NP-40, 1 mM Dithiothreitol (DTT)); bead binding buffer was the same minus DTT.

In a 384-well plate, 5 µL of RomA-6xHis (Query; WT or mutant form; concentration as noted) was combined with 5 µL of biotinylated nucleosome (Target; 10 nM final) in assay buffer and incubated for 30 mins at room temperature. A 10 µL mix of 2.5 µg/mL Ni-NTA acceptor beads (Table S8) and 10 µg/mL streptavidin donor beads (Table S8) in bead binding buffer was added and the plate incubated at room temperature for 60 mins. Alpha signal was measured on a *PerkinElmer 2104 EnVision* (680 nm laser excitation, 570 nm emission filter ± 50 nm bandwidth). Each reaction was performed in triplicate. Binding curves [Query : Target] were generated using a non-linear 4PL curve fit in GraphPad Prism [24], with $EC_{50}^{rel}$ values computed for specific comparisons. Where necessary, values beyond the hook point (indicating bead saturation / competition with unbound Query) were excluded and top signal constrained to average max signal for Target. In cases where signal never plateaued, signal was constrained to the average max signal within the assay.

**Captify™ luminex binding assays**

Assays to examine the interaction of antibodies (the Queries) with biotinylated PTM-defined nucleosomes (the Targets) in multiplex Luminex panels were performed in 96-well plates (Table S8) with assay buffer (20 mM HEPES pH 7.5, 150 mM NaCl, 0.01% Tween-20, 0.01% BSA). All bead handling was performed under subdued lighting, with continuous mixing to ensure monodispersion during incubations, and magnetic capture for wash steps.

Avidin-conjugated MagPlex® beads (*Luminex, Diasorin*) with spectrally distinct regions were used to assemble a 20-plex panel with fully defined biotinylated nucleosomes (Table S1 and Figure S1) as previously [28, 29, 30]. Balance, nucleosome integrity and identity (Nucleosome : bead region) within the panel were confirmed using anti-dsDNA (Table S10; 1/5, 1/50 and 1/500), anti-histone (Table S10), anti-H3.1/2 (Table S10) and an appropriate anti-PTM if available (Table S10). For each analysis, 50 µL multiplexed bead panel (20,000 beads/mL/region; 1,000 beads/region) was combined with 50 µL antibody (1:250, 1:1000 and 1:4000 from ~1 mg/ml unless otherwise specified). The reaction plate was incubated for 60 mins with shaking (800 rpm) before beads were washed for three cycles on a magnet using 100 µL assay buffer, shaking for two mins between each cycle. 100 µL of rabbit anti-IgG PE (1:100; Table S10) was added to each well and incubated for 30 mins with shaking. Beads were washed for two cycles, resuspended in 100 µL assay buffer, and Median fluorescence intensity (MFI) measured using the FLEXMAP-3D System (*Diasorin*) with a minimum of 50 events/region. Bar graphs were generated using GraphPad Prism Version 10.2.0 [24]. Each analysis was performed a minimum of twice with representative data shown.

**X-ray crystallography**

Protein crystallization was conducted using LegAS4 and RomA constructs that lacked residues 1 – 70 (Δ1-70) comprising their nuclear localization signal (NLS) and the linker between the NLS and the SET domain, which are predicted to be intrinsically disordered. Crystals of LegAS4 Δ1-70 construct bound to AdoMet or the product S-adenosylhomocysteine (AdoHcy) and $H3.1_{[1-21]}$ peptides were grown using the hanging-drop vapor diffusion method at 20 °C (Tables S2, S3, S4, S8). LegAS4 Δ1-70 (10mg/ml) was mixed with 1.5 mM AdoMet or AdoHcy and 1.5 mM peptide $H3.1_{[1-21]}$ in a solution containing 20 mM Tris pH 8.0, 100 mM NaCl, and 1 mM tris (2-carboxyethyl) phosphine (TCEP) (Table S8). The protein and ligand solutions were combined with an equal volume of crystallization solutions (100 mM Bis-Tris (pH 5.8, 6.0, 6.2, and 6.4), 10% – 20% (w/v) PEG 1500, and 30% (v/v) ethylene glycol). Crystals of the RomA Δ1-70/AdoMet/$H3_{[7-21]}$K14-Nle peptide complex were grown by hanging drop vapor diffusion as described for LegAS4 Δ1-70 using crystallization solutions composed of 100 mM sodium citrate pH 5.6, 10% –15% (w/v) PEG 4000, and 4-7% (v/v) isopropanol. Crystals of the LegAS4 ternary complexes were harvested and directly flash-frozen in liquid nitrogen, whereas the RomA crystals were harvested in their crystallization solution supplemented with 15% PEG 400 as the cryoprotectant (v/v) and then frozen in liquid nitrogen for X-ray diffraction screening and data collection.

X-ray diffraction data were collected under cryogenic conditions at 100 K at the Life Sciences Collaborative Access Team (LS-CAT), Advanced Photon Source (APS) at Argonne National Laboratory using the 21-ID-F or 21-ID-G beamlines. Diffraction data were indexed, integrated, and scaled using HKL2000 [31]. LegAS4 and RomA structures were determined by molecular replacement using the PHASER [32] in the Phenix software package with a previously determined structure of LegAS4 bound to AdoMet (PDB 5CZY) as the search model. Model building was conducted using COOT [33, 34], and structure refinement and validation were performed utilizing Phenix [35].

All structural figures and movies were generated using PyMOL (Schrödinger, Inc.). A simulated-annealing omit map for the histone $H3_{[1-21]}$ peptide and AdoHcy was generated for the structure of the LegAS4 Δ1-70/AdoHcy/$H3_{[1-21]}$ peptide complex using Phenix with the ligands omitted. The electrostatic surface potential for the structure of the LegAS4 Δ1-70/AdoHcy/$H3_{[1-21]}$ peptide complex was calculated using the Adaptive Poisson-Boltzmann Solver (APBS) in PyMOL [36, 37]. The modeling of H3K4me3 into the H3K4 binding pocket in the structure of the LegAS4 Δ1-70/AdoHcy/$H3_{[1-21]}$K4me3 peptide complex was accomplished by adding two methyl groups to the

$\varepsilon$-amino group of $H3_{[1-21]}$K4me1 using the Builder tool in PyMOL. The movie illustrating the binding of the $H3_{[1-21]}$K14-Nle peptide to LegAS4 (Supplemental Movie S1) was rendered using the coordinates of the LegAS4 $\Delta$1-70/AdoMet/$H3_{[1-21]}$K14-Nle peptide and LegAS4/AdoMet (PDB 5CZY) complexes using PyMOL.

**Sequence alignments**

The sequences of the N-terminal tails of histone H3 from representative protozoan and metazoan host species of *L. pneumophila* were aligned using Clustal Omega with Lasergene v18. The Histone H3 accession IDs used to generate the alignment include *Caenorhabditis elegans* (UniProt P08898), *Homo sapiens* (UniProt P68431), *Dictyostelium discoideum* (NCBI XP_646792.1), *Acanthamoeba castellanii* (NCBI XP_004342027.1), *Tetrahymena thermophila* (UniProt P69150), and *Naegleria gruberi* (NCBI EFC36852.1). In the sequence alignment, the backgrounds of the amino acids were colored according to their respective side chain properties using Adobe Illustrator. Invariant residues and amino acids that are conserved or semi-conserved are denoted under the sequence alignment.

Pairwise sequence alignment of the LpHKMT orthologues LegAS4 (UniProt Q5ZUS4) and RomA (NCBI: CAH12835.1) [38] from the *L. pneumophila* Philadelphia-1 and Paris strains, respectively, was performed with Clustal Omega with Lasergene v18 and rendered in Adobe Illustrator. The initial and new translation start sites (ITSS and NTSS) reported for LegAS4 are labeled at its N-terminus as described[39]. The background colors of the amino acids indicate the domain or subdomain in which they reside and were rendered using Adobe Illustrator. Residues that engage in hydrogen bonding with substrates or products are highlighted. Amino acids that are identical, conserved, or semi-conserved are denoted below the sequence alignment.

**SUPPLEMENTARY TABLES:**

**Table S1: Histone octamers and nucleosomes.**

| Octamers | Manufacturer | Catalog |
|---|---|---|
| H3.1 C110A *Xenopus laevis* octamers | The Histone Source | XOCT_H3C110A_10mg |
| H3.1K14M C110A *X. laevis* octamers | The Histone Source and this work | Custom order |
| H3.1R2A C110A *X. laevis* octamers | This work | N/A |
| H3.1K4A C110A *X. laevis* octamers | This work | N/A |
| **Nucleosome** | **Manufacturer** | **Catalog** |
| H3.1 C110A *X. laevis* | This work | N/A |
| ([H3.1R2A]$_2$) C110A *X. laevis* | This work | N/A |
| ([H3.1K4A]$_2$) C110A *X. laevis* | This work | N/A |
| ([H3.1K14M]$_2$) C110A *X. laevis* | This work | N/A |
| ([H2AR3me1]$_2$) | EpiCypher | 16-0359 |
| ([H2AR3me2a]$_2$) | EpiCypher | 16-0360 |
| ([H2AR3me2s]$_2$) | EpiCypher | 16-0361 |
| H2A tetra ac (H2AK5acK9acK13acK15ac]$_2$) | EpiCypher | 16-0376 |
| ([H2AK119Ub]$_2$) | EpiCypher | 16-0395 |
| ([H2BK120Ub]$_2$) | EpiCypher | 16-0396 |
| ([H2AXS139ph]$_2$) | EpiCypher | 16-0366 |
| ([H2AX]$_2$) | EpiCypher | 16-0013 |
| ([H2AZ.1]$_2$) | EpiCypher | 16-0014 |
| ([H2AZ.2]$_2$) | EpiCypher | 16-0015 |
| rNuc (all major histones) ([H3.1]$_2$) | EpiCypher | 16-0006 |
| ([H3.3]$_2$) | EpiCypher | 16-0011 |
| ([H3ΔN2]$_2$) | EpiCypher | 16-0023 |
| ([H3ΔN32]$_2$) | EpiCypher | 16-0016 |
| ([H3K4me2]$_2$), non-biotinylated DNA | EpiCypher | 16-1334 |
| ([H3K4me3]$_2$), non-biotinylated DNA | EpiCypher | 16-1316 |
| ([H3K4me1]$_2$) | EpiCypher | 16-0321 |
| ([H3K4me2]$_2$) | EpiCypher | 16-0334 |
| ([H3K4me3]$_2$) | EpiCypher | 16-0316 |

| | | |
|---|---|---|
| ([H3K9me1]$_2$) | EpiCypher | 16-0325 |
| ([H3K9me2]$_2$) | EpiCypher | 16-0324 |
| ([H3K9me3]$_2$) | EpiCypher | 16-0315 |
| ([H3K14un]$_2$) [versaNuc, WT] (Figure S1) | EpiCypher | N/A |
| ([H3K14me1]$_2$) [versaNuc] (Figure S1) | EpiCypher | N/A |
| ([H3K14me2]$_2$) [versaNuc] (Figure S1) | EpiCypher | N/A |
| ([H3K14me3]$_2$) [versaNuc] (Figure S1) | EpiCypher | N/A |
| ([H3K27me1]$_2$) | EpiCypher | 16-0338 |
| ([H3K27me2]$_2$) | EpiCypher | 16-0339 |
| ([H3K27me3]$_2$) | EpiCypher | 16-0317 |
| ([H3K36me1]$_2$) | EpiCypher | 16-0322 |
| ([H3K36me2]$_2$) | EpiCypher | 16-0319 |
| ([H3K36me3]$_2$) | EpiCypher | 16-0320 |
| ([H3K79me1]$_2$) | EpiCypher | 16-0367 |
| ([H3K79me2]$_2$) | EpiCypher | 16-0368 |
| ([H3K79me3]$_2$) | EpiCypher | 16-0369 |
| ([H3K4ac]$_2$) | EpiCypher | 16-0342 |
| ([H3K9ac]$_2$) | EpiCypher | 16-0314 |
| ([H3K9bu]$_2$) | EpiCypher | 16-0371 |
| ([K3K9cr]$_2$) | EpiCypher | 16-0351 |
| ([H3K14ac]$_2$) | EpiCypher | 16-0343 |
| ([H3K18ac]$_2$) | EpiCypher | 16-0372 |
| ([H3K18bu]$_2$) | EpiCypher | 16-0373 |
| ([H3K18cr]$_2$) | EpiCypher | 16-0337 |
| ([H3K23ac]$_2$) | EpiCypher | 16-0364 |
| ([H3K27ac]$_2$) | EpiCypher | 16-0365 |
| ([H3K27bu]$_2$) | EpiCypher | 16-0384 |
| ([H3K27cr]$_2$) | EpiCypher | 16-0383 |
| ([H3K36ac]$_2$) | EpiCypher | 16-0378 |
| ([H3R2me1]$_2$) | EpiCypher | 16-0340 |
| ([H3R2me2a]$_2$) | EpiCypher | 16-0341 |
| ([H3R2me2s]$_2$) | EpiCypher | 16-0355 |
| ([H3R8me1]$_2$) | EpiCypher | 16-0340 |

| | | |
|---|---|---|
| ([H3R8m2a]$_2$) | EpiCypher | 16-0341 |
| ([H3R8me2s]$_2$) | EpiCypher | 16-0355 |
| ([H3R17me1]$_2$) | EpiCypher | 16-0382 |
| ([H3R17me2a]$_2$) | EpiCypher | 16-0375 |
| ([H3S10ph]$_2$) | EpiCypher | 16-0345 |
| ([H4 ΔN15]$_2$) | EpiCypher | 16-0018 |
| ([H4K20me1]$_2$) | EpiCypher | 16-0331 |
| ([H4K20me2]$_2$) | EpiCypher | 16-0332 |
| ([H4K20me3]$_2$) | EpiCypher | 16-0333 |
| ([H4K5ac]$_2$) | EpiCypher | 16-0352 |
| ([H4K8ac]$_2$) | EpiCypher | 16-0353 |
| ([H4K12ac]$_2$) | EpiCypher | 16-0312 |
| ([H4K16ac]$_2$) | EpiCypher | 16-0354 |
| ([H4K20ac]$_2$) | EpiCypher | 16-0377 |
| H4 tetra ac ([H4K5acK8acK12acK16ac]$_2$) | EpiCypher | 16-0313 |
| ([H4R3me1]$_2$) | EpiCypher | 16-0356 |
| ([H4R3me2a]$_2$) | EpiCypher | 16-0357 |
| ([H4R3me2s]$_2$) | EpiCypher | 16-0358 |

**Table S2: LpHKMT constructs.**

| LpHKMT Construct | Residue Range | Application |
|---|---|---|
| RomA (full length) | 1 – 532 | EMSAs, methyltransferase assays, Captify-Alpha |
| RomA ΔNLS | 50 – 532 | EMSAs, Captify-Alpha |
| RomA Δ1-70 | 71 – 532 | Crystallography, ITC |
| LegAS4 (full length) | 1 – 532 | EMSAs, methyltransferase assays |
| LegAS4 Δ1-70 | 71 – 532 | Crystallography |

### Table S3: Crystallographic and refinement data.

| | LegAS4•AdoHcy• H3$_{[1-21]}$ | LegAS4•AdoMet• H3$_{[1-21]}$K14-Nle | LegAS4•AdoHcy• H3$_{[1-21]}$K4me1 | LegAS4•AdoHcy• H3$_{[1-21]}$R2me1 | LegAS4•AdoHcy• H3$_{[1-21]}$R2me2a | LegAS4•AdoHcy• H3$_{[1-21]}$R2me2s | RomA•AdoMet• H3$_{[7-21]}$K14-Nle |
|---|---|---|---|---|---|---|---|
| **Beamline (APS)** | 21-ID-F | 21-ID-F | 21-ID-G | 21-ID-G | 21-ID-G | 21-ID-G | 21-ID-F |
| **Data Collection** | | | | | | | |
| Wavelength (Å) | 0.97872 | 0.97872 | 0.97857 | 0.97857 | 0.97857 | 0.97857 | 0.97872 |
| Space group | $P3_121$ | $P3_121$ | $P3_121$ | $P3_121$ | $P3_121$ | $P3_121$ | $P12_11$ |
| Cell dimensions | | | | | | | |
| *a, b, c* (Å) | 68.53 68.53 198.87 | 68.42 68.42 198.22 | 68.59 68.59 198.88 | 68.55 68.55 201.28 | 68.72 68.72 198.74 | 68.45 68.45 198.96 | 62.262 65.16 123.25 |
| *α, β, γ* (°) | 90 90 120 | 90 90 120 | 90 90 120 | 90 90 120 | 90 90 120 | 90 90 120 | 90 100.73 90 |
| Resolution (Å) | 33.16 - 1.60 (1.63 - 1.6)[1] | 38.02 - 1.81 (1.83 - 1.80)[1] | 44.24 - 1.75 (1.78 - 1.75)[1] | 33.79 - 1.60 (1.63 – 1.60)[1] | 33.86 - 1.80 (1.83 – 1.8)[1] | 33.73 - 1.75 (1.78 - 1.75)[1] | 40.37 - 1.75 (1.77 - 1.75)[1] |
| $R_{merge}$ | 0.057 (1.171) | 0.124 (1.448) | 0.067 (1.325) | 0.051 (0.635) | 0.055 (1.513) | 0.072 (1.349) | 0.081 (0.840) |
| CC1/2 | (0.558) | (0.513) | (0.621) | (0.803) | (0.527) | (0.596) | (0.642) |
| $\langle I/\sigma I \rangle$ | 18.7 (1.46) | 12.0 (1.70) | 18.2 (1.49) | 22.5 (2.62) | 11.7 (1.16) | 12.5 (1.62) | 8.41 (2.35) |
| Completeness (%) | 98.5 (83.2) | 99.6 (95.2) | 99.5 (98.0) | 98.1 (81.8) | 96.1 (88.5) | 94.3 (96.3) | 92.5 (80.5) |
| Redundancy | 10.2 (5.9) | 8.7 (6.8) | 10.7 (8.6) | 10.3 (6.6) | 6.3 (5.0) | 6.6 (6.7) | 3.7 (2.7) |
| **Refinement** | | | | | | | |
| Resolution | 33.16 - 1.60 | 38.02 - 1.81 | 44.24 - 1.75 | 33.79 - 1.60 | 33.86 - 1.80 | 33.73 – 1.75 | 40.37 - 1.75 |
| No. of Reflections | 71,389 | 50,120 | 55,431 | 71,886 | 49,666 | 52,148 | 90,137 |
| $R_{work}/R_{free}$ | 0.154/0.187 | 0.165/0.208 | 0.181/0.204 | 0.153/0.191 | 0.196/0.228 | 0.183/0.207 | 0.210/0.248 |
| No. of Non-Hydrogen Atoms | 4245 | 4282 | 4257 | 4267 | 4013 | 4159 | 7998 |
| Protein | 3811 | 3716 | 3764 | 3755 | 3704 | 3690 | 7015 |
| Ligand | 85 | 127 | 141 | 162 | 81 | 179 | 54 |
| Water | 349 | 439 | 352 | 349 | 228 | 289 | 929 |
| B-factors | 32.62 | 28.56 | 33.67 | 27.59 | 46.64 | 31.62 | 29.24 |
| Protein | 33.86 | 26.82 | 32.70 | 26.11 | 46.62 | 30.84 | 28.38 |
| Ligand/ion | 35.54 | 34.86 | 39.38 | 37.80 | 44.77 | 37.64 | 19.79 |
| Water | 42.35 | 41.42 | 41.73 | 38.75 | 47.58 | 37.84 | 36.31 |
| R.M.S. Deviations | | | | | | | |
| Bond lengths (Å) | 0.013 | 0.014 | 0.004 | 0.012 | 0.003 | 0.006 | 0.006 |
| Bond angles (°) | 1.200 | 1.190 | 0.640 | 1.17 | 0.58 | 0.76 | 0.76 |
| **Ramachandran Plot** | | | | | | | |
| Favored (%) | 98.92 | 98.91 | 98.70 | 99.14 | 98.92 | 99.35 | 98.63 |
| Allowed (%) | 1.08 | 1.09 | 1.30 | 0.86 | 1.08 | 0.65 | 1.37 |
| Outliers (%) | 0 | 0 | 0 | 0 | 0 | 0 | 0 |

[1] Reported values for the highest-resolution shell are shown in parentheses.

**Table S4: Histone H3 peptides.**

| Manufacturer | Amino Acid Sequence | Peptide Name |
|---|---|---|
| Biosynth | NH2- $A_1$RTKQTARKSTGGKAPRKQL$A_{21}$ - NH2 | [H3.1$_{[1-21]}$] |
| Biosynth | NH2-$A_1$RTKQTARKSTGG-(Nle14)-APRKQL$A_{21}$- NH2 | [H3.1$_{[1-21]}$K14-Nle] |
| Biosynth | NH2- $A_1$RTK$4_{me1}$QTARKSTGGKAPRKQL$A_{21}$ - NH2 | [H3.1$_{[1-21]}$K4me1] |
| Biosynth | NH2- $A_1$R$2_{me1}$TKQTARKSTGGKAPRKQL$A_{21}$ - NH2 | [H3.1$_{[1-21]}$R2me1] |
| Biosynth | NH2- $A_1$R$2_{me2a}$TKQTARKSTGGKAPRKQL$A_{21}$ - NH2 | [H3.1$_{[1-21]}$R2me2a] |
| Biosynth | NH2- $A_1$R$2_{me2s}$TKQTARKSTGGKAPRKQL$A_{21}$ - NH2 | [H3.1$_{[1-21]}$R2me2s] |
| Biosynth | Ac-$A_7$RKSTGG-(Nle14)-APRKQL$A_{21}$-NH2 | [H3.1$_{[7-21]}$K14-Nle] |

**Table S5: LegAS4-histone H3 hydrogen bonding.**

| Hydrogen Bonding Between Histone H3 and LegAS4 | | | | | | |
|---|---|---|---|---|---|---|
| **Histone H3** | **H3 Atom Name** | **Group Type** | **Distance (Å)** | **LegAS4 Residue** | **LegAS4 Atom Name** | **Group Type** |
| H3A1 | N | Main chain | 2.8 | Asp184 | OD2 | Side chain |
| H3R2 | NE | Side chain | 3.1 | Asp438 | OD1 | Side chain |
| H3R2 | NH1 | Side chain | 3.3 | Tyr223 | OH | Side chain |
| H3R2 | NH2 | Side chain | 2.7 | Asp438 | OD1 | Side chain |
| H3R2 | O | Main chain | 3.0 | Ile434 | N | Main chain |
| H3K4 | N | Main chain | 2.9 | Thr432 | O | Main chain |
| H3K4 | NZ | Side chain | 3.0 | Phe431 | O | Main chain |
| H3K4 | NZ | Side chain | 2.8 | Asp440 | OD2 | Side chain |
| H3K4 | NZ | Side chain | 2.8 | Asn480 | ND2 | Side chain |
| H3Q5 | OE1 | Side chain | 2.8 | Thr432 | OG1 | Side chain |
| H3K9 | NZ | Side chain | 2.7 | Asn221 | O | Main chain |
| H3T11 | O | Main chain | 3.2 | Lys157 | NZ | Side chain |
| H3G12 | N | Main chain | 3.0 | Tyr223 | O | Main chain |
| H3G13 | O | Main chain | 2.7 | Thr222 | OG1 | Side chain |
| H3K14 | N | Main chain | 2.8 | Ala160 | O | Main chain |
| H3K14 | NZ | Side chain | 2.9 | Tyr135 | OH | Side chain |
| H3K14 | O | Main chain | 2.8 | Thr162 | N | Main chain |
| H3A15 | N | Main chain | 2.9 | Tyr220 | O | Main chain |
| H3A15 | O | Main chain | 2.9 | Asn221 | ND2 | Side chain |
| H3R17 | NH1`A | Side chain | 2.7 | Glu190 | OE1 | Side chain |
| H3R17 | NH2`A | Side chain | 3.0 | Asp187 | OD2 | Side chain |
| H3R17 | NH1`B | Side chain | 2.9 | Asn221 | OD1 | Side chain |
| H3R17 | NH2`B | Side chain | 2.9 | Asp187 | OD2 | Side chain |

**Table S6: Strains.**

| Strain | Relevant Properties | Source |
|---|---|---|
| *E. coli* | | |
| Rosetta2-DE3 | Derivative of BL21 strain, optimized for expressing 7 tRNA codons. | Novagen-71397 |
| DH5α | Used for cloning in preparation for protein expression and further analysis by: EMSA, Alpha, and MTase-Glo. | Invitrogen-18258012 |
| DY330* | Electrocompetent $\lambda$ (lambda) red recombination. | [5] |
| *L. pneumophila* | | |
| MB534* | Lp02 Δ*flaA::kan* mutant (from Lp02 wild type; Δ*thyA* Δ*hsdR* Δ*rpsL*) | [1] |
| TK55 | Lp02 Δ*flaA::kan* Δ*legAS4* mutant* | This work |
| TK56 | Lp02 Δ*flaA::kan* Δ*legAS4* + pSU-pZLCm* | This work |
| TK57 | Lp02 Δ*flaA::kan* Δ*legAS4* + pSU-pZLCm-*legAS4** | This work |
| TK58 | Lp02 Δ*flaA::kan* Δ*legAS4* + pSU-pZLCm-*romA** | This work |
| TK59 | Lp02 Δ*flaA::kan* Δ*legAS4* + pSU-pZLCm-*legAS4*-CD* | This work |

*Integrated between *lpg2528* and *lpg2529* (considered a neutral site) in Lp02 chromosome[7].

### Table S7: Primers.

| Num. | Name | Sequence 5' – 3' | Primer Use |
|---|---|---|---|
| **Construction of *legAS4* deletion mutant** | | | |
| 192 | Lpg1718Fwd | AAACTATATCGCCTAGCTGGACACCGGG | Amplify *lpg1718* + ~500bp of flanking DNA |
| 193 | Lpg1718Rev | AAACACTTTGCCGATTCGAAGCAATTAGC | |
| 185 | Lpg1718_P0_F | GAGTATAATGGCTCCTTAATGATTATTTTATATGATTGTGTAGGCTGGAGCTGCTTC | Amplify *cat* cassette with homology to *lpg1718* |
| 186 | Lpg1718_P2_R | TAAGATATGTAAACAGGAAATATCAAATCAAAAAAACATATGAATATCCTCCTTAGTTCC | |
| | **Construction of complementation vectors** | | |
| 268 | GibsonApKD3 | CGACGTTGTAAAACGACGGCCAGTGAATTCGAGCTCTGTAGGCTGGAGCTGCTTCG | Amplify Gibson insert from pKD3 |
| 270 | GibsonBpKD3 | CAGGTCGACTCTAGAGGATCCCCGGGTACCGAGCTCAGCCCTTAGAGCCTCTCAAAG C | |
| 269 | GibsonApSUpZLKm | CGAAGCAGCTCCAGCCTACAGAGCTCGAATTCACTGGCCGTCGTTTTACAACGTCG | Amplify Gibson backbone from pSU-pZLKm |
| 271 | GibsonBpSUpZLKm | GCTTTGAGAGGCTCTAAGGGCTGAGCTCGGTACCCGGGGATCCTCTAGAGTCGACCT G | |
| 127bis | LegAS4Fbis | CGTGGCAGGGTACCCTATATCGCCTAGCTGGACACCGGG | Amplify *legAS4* + ~500bp flanking DNA adding KpnI and BamHI |
| 128 | LegAS4Rev | CGTGGCAGGGATCCCACTTTGCCGATTCGAAGCAATTAGC | |
| 129bis | ParisF | CGTGGCAGGGTACCTCATGAAATACACGGCCAATTCCATGTCC | Amplify *legAS4* + ~500bp flanking DNA adding KpnI and BamHI |
| 128 | LegAS4Rev | CGTGGCAGGGATCCCACTTTGCCGATTCGAAGCAATTAGC | |
| | **Construction of catalytically deficient *legAS4* mutant** | | |
| 202 | Y148F_Fwd | GTATTGGTATTTTCACAGGAGAAGTATATTCTGAACAAGAATTTGAAC | Site-directed point mutagenesis PCR |
| 203 | Y148F_RED | CTCCTGTGAAAATACCAATACAAGTCCCCTTTGGGATATCTTCCCTGGC | |
| 204 | Y172F_FWD | CTGATAAAAGTTTCGCGATGTATGTTGGGGGACGGGTAATTGATGCAGC | |
| 205 | Y172F_REV | ACATCGCGAAACTTTTATCAGAACCAACATGTTCCTTTAGATATTGTTC | |
| 206 | Y233F_FWD | CTATTAATCAACTTTAATACCTACGAAGAGCAGGCCTCCAGGTATTATTAC | |

| | | | |
|---|---|---|---|
| 207 | Y233F_REV | AGGTATTAAAGTTGATTAATAGTTGTTGTCCTGCCTTGATGTTTTTAGTGG | |
| 246 | N194A_FWD | CTCGTTATATCGCTTTTTCGGATAGTCAGGATAATGCAGAGTTTGTTG | |
| 247 | N194A_REV | CCGAAAAAGCGATATAACGAGTCAAATTACCTTTTCTTGCTGCATCAATTACC | |
| 284 | S171A_FWD | GTTCTGATAAAGCTTTCGCGATGTATGTTGGGGGACGGGTAATTGATG | |
| 285 | S171A_REV | TCGCGAAAGCTTTATCAGAACCAACATGTTCCTTTAGATATTGTTC | |
| 286 | Y244F_FWD | GCCTCCAGGTATTTTTACTTTCTTAATCCTGGCGATGGTTGGCTATCGG | |
| 287 | Y244F_REV | GAAAGTAAAAATACCTGGAGGCCTGCTCTTCGTAGGTATTAAAGTTG | |
| | **HA tag insertions** | | |
| 260 | HAtagCterm_legAS4_F | CTTGAAAGCAAGTTTTTTTATCCATATGATGTTCCAGATTATGCTTGATTTGATATTTCCTGTTTACATATC | *legAS4* site-directed insertion mutagenesis PCR |
| 261 | HAtagCterm_legAS4_R | ATAAAAAAACTTGCTTTCAAGTGTGATACTTGAATCCATAATACCTAATTG | |
| 262 | HAtagCterm_romA_F | CTCGAAAGCAAGTTTTTTTATCCATATGATGTTCCAGATTATGCTTGATTTGATATTTCCTGTTTACATATC | *romA* site-directed mutagenesis PCR |
| 263 | HAtagCterm_romA_R | ATAAAAAAACTTGCTTTCGAGTGTGATATTTGAATCCATAATACCTAATTG | |
| | **Construction of *RomA* mutants with N-terminal 6xHis-SUMO** | | |
| | K157A_FWD | GGGTAGCGATGCAAGCTATGCAATGTATGTTGGTGGTCGCGTTGTTG | Site-directed point mutagenesis PCR |
| | K157A_REV | CATAGCTTGCATCGCTACCCACATGTTCCATCAGATACTGTTCAAAC | |
| | D184A_FWD | TAACTTTAGCGCTAGCCAGGATAATGCCGAATTTGTTGAAACCACACTG | |
| | D184A_REV | CCTGGCTAGCGCTAAAGTTAATATAGCGGGTCAGATTACCTTTACGTG | |
| | N221A_FWD | GATCAACTATGCCACCTATGAAGAACAGGCAAGCCGCTATTATTACTTC | |
| | N221A_REV | CATAGGTGGCATAGTTGATCAGCAGTTGCTGACCTGCTTTAATGTTTTTG | |
| | T222A_FWD | CAACTATAACGCCTATGAAGAACAGGCAAGCCGCTATTATTACTTCTG | |
| | T222A_REV | CTTCATAGGCGTTATAGTTGATCAGCAGTTGCTGACCTGCTTTAATG | |
| | Y223A_FWD | CTATAACACCGCTGAAGAACAGGCAAGCCGCTATTATTACTTTCTGAATCC | |
| | Y223A_REV | CTGTTCTTCAGCGGTGTTATAGTTHATCAHCAHTTHCTHACCTHCTTTAATG | |
| | Y223F_FWD | CTATAACACCTTTGAAGAACAGGCAAGCCGCTATTATTACTTTCTGAATCC | |
| | Y223F_REV | CTGTTCTTCAAAGGTGTTATAGTTGATCAGCAGTTGCTGACCTGCTTTAATG | |
| | T432A_FWD | CGAATATTTCGCCTACATCGACGAGAACGATTTCGATATTGTGATGCAC | |
| | T432A_REV | CGATGTAGGCGAAATATTCGTTGATATCGATCGGATTCTGGCCAATCAG | |
| | D438A_FWD | CGACGAGAACGCTTTCGATATTGTGATGCACTGCTACAACAACAAACTG | |
| | D438A_REV | CAATATCGAAAGCGTTCTCGTCGATGTAGGTGAAATATTCGTTGATATCGATC | |

| | | | |
|---|---|---|---|
| | D438N_FWD | CGACGAGAACAATTTCGATATTGTGATGCACTGCTACAACAACAAACTG | |
| | D438N_REV | CAATATCGAAATTGTTCTCGTCGATGTAGGTGAAATATTCGTTGATATCGATC | |
| | D440A_FWD | GAACGATTTCGCTATTGTGATGCACTGCTACAACAACAAACTGTTTGATAAAG | |
| | D440A_REV | GCATCACAATAGCGAAATCGTTCTCGTCGATGTAGGTGAAATATTCGTTGA | |
| | D440N_FWD | GAACGATTTCAATATTGTGATGCACTGCTACAACAACAAACTGTTTGATAAAG | |
| | D440N_REV | GCATCACAATATTGAAATCGTTCTCGTCGATGTAGGTGAAATATTCGTTGA | |
| | N480A_FWD | CAACCAGTTTGCCATTAACGCATTTCGCAAAGCGATCAAGGATTTTAAC | |
| | N480A_REV | GCGTTAATGGCAAACTGGTTGTGGCCTTCATTATCGCTCATATAGTTGC | |
| | Y135F_FWD | GTATTGGTATTTTTACCGGTGAAGTTTATAGCGAGCAAGAGTTTGAACAG | |
| | Y135F_REV | CACCGGTAAAAATACCAATACAGGTGCCTTTCGGAATATCTTCACGTG | |
| | S158A & Y159F,_FWD | AGCGATAAAGCCTTTGCAATGTATGTTGGTGGTCGCGTTGTTGATGCAGC | |
| | S158A & Y159F,_REV | CATTGCAAAGGCTTTATCGCTACCCACATGTTCCATCAGATACTGTTC | |
| | N181A_FWD | CCCGCTATATTGCCTTTAGCGATAGCCAGGATAATGCCGAATTTGTTG | |
| | N181A_REV | CGCTAAAGGCAATATAGCGGGTCAGATTACCTTTACGTGCTGCATCAAC | |
| | Y220F_FWD | GCTGATCAACTTTAACACCTATGAAGAACAGGCAAGCCGCTATTATTAC | |
| | Y220F_REV | AGGTGTTAAAGTTGATCAGCAGTTGCTGACCTGCTTTAATGTTTTTGGTG | |
| | Y231F_FWD | GGCAAGCCGCTATTTTTACTTTCTGAATCCTGGTGATGGTTGGCTGAGC | |
| | Y231F_REV, Y220F_REV | GTAAAAATAGCGGCTTGCCTGTTCTTCATAGGTGTTAAAGTTGATCAGC | |
| | **Construction of in-house Nucleosomes** | | |
| | H3.1C110A_FWD | CCAACCTGGCCGCCATCCACGCCAAGAGGGTCACCATCATGCCCAAGG | H3.1 C110A |
| | H3.1C110A_REV | GCGTGGATGGCGGCCAGGTTGGTGTCCTCAAAGAGAGCGACCAGATAAGCC | |
| | H3.1K14M_FWD | CACCGGAGGGATGGCTCCCCGCAAGCAGCTGGCCACCAAGGCAGCCAG | H3.1K14M |
| | H3.1K14M_REV | CGGGGAGCCATCCCTCCGGTGGATTTACGGGCGGTCTGCTTGGTACGGGC | |
| | H3.1K4A_FWD | GGCCCGTACCGCGCAGACCGCCCGTAAATCCACCGGAGGGAAGGC | H3.1K4A |
| | H3.1K4A_REV | CCCGCCAGACGCGCCATGCCCGGTACCATACATATAGAGGAAGAATTTCAATTTG | |
| | H3.1R2A_FWD | GCCTTCCTCCGGTGGATTTACGGGCGGTCTGCTTGGTAGCGGCCATGG | H3.1R2A |
| | H3.1R2A_REV | CAGACGAACCATCGCCGGTACCATACATATAGAGGAAGAATTTCAATTTG | |
| | **Construction of 6xHis C-terminal on Strep-SUMO-RomA constructs for Alpha** | | |
| | RomA stop-codon to Alanine_FWD | CCAAATTCTTTGCACTCGAGCACCACCACCACCACCACTGAGATCCGGCTGC | |

| | | |
|---|---|---|
| | RomA stop-codon to Alanine_REV | GGAGCTCGAGTGCAAAGAATTTGGATTCCAGGGTAATATTGCTATCCATAATG |
| | Δ1-49 RomA_FWD | GATTGGTGGATCCATGACCCATCTGCCTGGTAACATTTATACCCTGTTTAC |
| | Δ1-49 RomA_REV | CAGATGGGTCATGGATCCACCAATCTGTTCTCTGTGAGCCTCAATAATATC |
| | **Construction of LegAS4 ΔNLS** | |
| | LegAS4ΔNLS_FWD | GCACGTCGGGATCCGATGAATGGATCGATACCAGC |
| | LegAS4ΔNLS_REV | CGTGGCAGCTCGAGTTAGAAAAACTTGCTTTCCAGGGTAATGC |

**Table S8: Reagents.**

| Name | Company | Catalog |
|---|---|---|
| pGEM®-T Easy Vector System | Promega | A1360 |
| RPMI 1640 | Corning | 10-040-CM |
| Fetal Bovine Serum, heat inactivated | Invitrogen | A5256801 |
| GlutaMax | Invitrogen | 35050061 |
| Human M-CSF lyophilized powder | R&D Systems | 216-MC-025 |
| Penicillin-Streptomycin (10K U/mL) | Invitrogen | 15140122 |
| 2-Mercaptoethanol (cell culture grade) | Fisher | 50-165-7065 |
| Gibco™ PBS, pH 7.4 ($Mg^{2+}$, $Ca^{2+}$, and Phenol Red free) | Fisher | 10010023 |
| HALT Protease inhibitor, EDTA-free | Thermo Scientific | 78425 |
| Protease inhibitor cocktail | Sigma-Aldrich | P8465 |
| Pierce™ BCA Protein Assay Kit | Thermo Scientific | 23225 |
| Laemmli Buffer | Bio-Rad | 161-0747 |
| 4-20% precast polyacrylamide gels | Bio-Rad | 4561096 |
| Poly(vinylidene difluoride) (PVDF) 0.45 μm pore | Millipore | IPVH15150 |
| Pierce™ ECL Western Blotting Substrate | Thermo Scientific | 32109 |
| Histone H3.1 Human, Recombinant | NEB | M2503 |
| S-(5'-Adenosyl)-L-methionine (AdoMet) | Cayman Chemical | 13956 |
| TALON® Superflow™ histidine-tagged resin | Cytiva | 28957502 |
| HisPur™ Ni-NTA Superflow Agarose | Thermo Scientific | 25215 |
| Strep-Tactin Superflow Plus resin | Qiagen | 30004 |
| Recombinant Albumin | NEB | B9200S |
| 5% Tris-Borate-EDTA (TBE) precast gels | Bio-Rad | 4565016 |
| 10,000X SYBR Gold Gel Stain in DMSO | Invitrogen | S11494 |
| MTase-Glo™ assay | Promega | V7601 |
| 384-well ProxiPlate | Revvity | 6008280 |
| S-(5'-Adenosyl)-L-homocysteine (AdoHcy) | Sigma-Aldrich | A9384 |
| Plasmid Giga-prep | Qiagen | 12191 |
| *Eco*RV-HF restriction endonuclease | NEB | R3195S |
| Source 15Q Strong Anion Exchange resin | Cytiva | 17094705 |
| Superdex 200 10/300 GL | Cytiva | 28-9909-44 |
| 2K MWCO Slide-A-Lyzer™ | Thermo Scientific | 66212 |

| | | |
|---|---|---|
| 3.5K MWCO Slide-A-Lyzer™ | Thermo Scientific | 66330 |
| 3.5 kDa SnakeSkin™ dialysis tubing | Thermo Scientific | 88242 |
| 10K MWCO Slide-A-Lyzer™ | Thermo Scientific | 66380 |
| 50K MWCO Vivaspin® 500 spin column | Cytiva | 28932236 |
| NaCl | Fisher | BP358 |
| KCl | Sigma-Aldrich | P3911 |
| EDTA | Sigma-Aldrich | E6758 |
| Imidazole | Fisher | 03196 |
| Sodium monophosphate | Sigma-Aldrich | S0751 |
| Sodium citrate dihydrate | Fisher | S279 |
| TRIS | Thermo Scientific | 17926 |
| Bis-TRIS | Fisher | BP301 |
| DTT (dithiothreitol) | Thermo Scientific | R0861 |
| IPTG (Isopropyl-B-D-thiogalatopyranoside) | IBI Scientific | IB02125 |
| TCEP (Tris (2-Carboxyethyl) phosphine Hydrochloride) | GoldBio | TCEP1 |
| PEG 400 | Sigma-Aldrich | PX1286B |
| PEG 1500 | Thermo Scientific | A16241 |
| PEG 4000 | Thermo Scientific | A16151 |
| Isopropanol | Sigma-Aldrich | I9030 |
| Ethylene glycol | Thermo Scientific | 146750010 |
| D-(+)-Glucose | Sigma-Aldrich | G5767 |
| Glycerol | Fisher | BP229 |
| Luria-Agar | RPI | L24022 |
| Terrific Broth | Fisher | BP2468 |
| LB Broth, Miller | Fisher | BP1426 |
| 2xYT Broth | RPI | X15600 |
| Kanamycin sulfate | Fisher | BP906 |
| Chloramphenicol | Sigma-Aldrich | C0378 |
| 24-well crystallization plates | Hampton Research | HR3-141 |
| 96-well crystallization plates | sptlabtech | 4150-05700 |
| Ni-NTA AlphaScreen Acceptor Beads | PerkinElmer | 6760619 |
| AlphaScreen Streptavidin Donor Beads | PerkinElmer | 6760002 |
| 96-well Luminex plates | GreinerBio | 655900 |

**Table S9. Plasmids.**

| Plasmid | Description | Source |
|---|---|---|
| *X. laevis* pET3d-Histone H2A | Expression plasmid encoding Histone H2A, AmpR | Uhn-Soo Cho |
| *X. laevis* pET3d-Histone H2B | Expression plasmid encoding Histone H2B, AmpR | Uhn-Soo Cho |
| *X. laevis* pET3d-Histone H3.1 | Expression plasmid encoding Histone H3.1, AmpR | Uhn-Soo Cho |
| *X. laevis* pET3d-Histone H4 | Expression plasmid encoding Histone H4, AmpR | Uhn-Soo Cho |
| pGEM-T Easy | Cloning vector, AmpR | Promega |
| pKD3 | Source of chloramphenicol gene flanked by Flp recognition targets, CamR | [4] |
| pBSFlp | Source of Flp recombinase, GentR | [6] |
| pGEM-*legAS4* | pGEM with *legAS4* and ~500 bp 3' and 5' flanking DNA, AmpR | This work |
| pGEM-*legAS4::cat* | pGEM with *cat*-disrupted *legAS4* and ~500 bp 3' and 5' flanking DNA, AmpR, CamR | This work |
| pSU-pZLKm | Cloning plasmid; template for construction of pSU-pZLCm; KanR | [7] |
| pSU-pZLCm | Cloning plasmid; vector for chromosomal integration into intergenic region of *lpg2528* and *lpg2529*; CamR | This work |
| pSU-pZLCm-legAS4 | Plasmid for insertion of *legAS4* with a C-terminal HA tag into Lp02 chromosome between *lpg2528* and *lpg2529* by homologous recombination; CamR | This work |
| pSU-pZLCm-romA | Plasmid for insertion of *romA* with a C-terminal HA tag into Lp02 chromosome between *lpg2528* and *lpg2529* by homologous recombination; CamR | This work |
| pSU-pZLCm-legAS4-CD | Plasmid for insertion of mutated *legAS4* with a C-terminal HA tag, producing a catalytically defective LegAS4, into Lp02 chromosome between *lpg2528* and *lpg2529* by homologous recombination; CamR | This work |

| | | |
|---|---|---|
| ppSUMO vector | Cloning vector; template containing N-terminal 6xHis-SUMO sequence for insertion of *romA* mutants and improvement in solubility; CamR | This work |
| 601-tandem repeats | 601 (147 bp) Widom DNA, with EcoRV restriction sites. | Uhn-Soo Cho |

**Table S10: Antibodies.**

| Target | Company | Cat. No. |
|---|---|---|
| β-actin | BioLegend | 643808 |
| H3 | Cell Signaling | 9715 |
| H3K14me2 | Invitrogen | MA-3059 |
| Anti-mouse HRP-conjugated | Millipore Sigma | AP308P |
| Anti-rabbit HRP-conjugated | Millipore Sigma | AP307P |
| Anti-dsDNA | EMD Millipore | MAB030 |
| Anti-Histone | Millipore Sigma | MAB3422 |
| Anti-Histone H3.1/3.2 | Active Motif | 61629 |
| Anti-Rabbit IgG | Biolegend | 406421 |
